# Neuronal functional connectivity is impaired in a layer dependent manner near the chronically implanted microelectrodes

**DOI:** 10.1101/2023.11.06.565852

**Authors:** Keying Chen, Adam Forrest, Guillermo Gonzalez Burgos, Takashi D.Y. Kozai

## Abstract

**Objective:** This study aims to reveal longitudinal changes in functional network connectivity within and across different brain structures near the chronically implanted microelectrode. While it is well established that the foreign-body response (FBR) contributes to the gradual decline of the signals recorded from brain implants over time, how does the FBR impact affect the functional stability of neural circuits near implanted Brain-Computer Interfaces (BCIs) remains unknown. This research aims to illuminate how the chronic FBR can alter local neural circuit function and the implications for BCI decoders.

**Approach:** This study utilized multisite Michigan-style microelectrodes that span all cortical layers and the hippocampal CA1 region to collect spontaneous and visually-evoked electrophysiological activity. Alterations in neuronal activity near the microelectrode were tested assessing cross-frequency synchronization of LFP and spike entrainment to LFP oscillatory activity throughout 16 weeks after microelectrode implantation.

**Main Results:** The study found that cortical layer 4, the input-receiving layer, maintained activity over the implantation time. However, layers 2/3 rapidly experienced severe impairment, leading to a loss of proper intralaminar connectivity in the downstream output layers 5/6. Furthermore, the impairment of interlaminar connectivity near the microelectrode was unidirectional, showing decreased connectivity from Layers 2/3 to Layers 5/6 but not the reverse direction. In the hippocampus, CA1 neurons gradually became unable to properly entrain to the surrounding LFP oscillations.

**Significance:** This study provides a detailed characterization of network connectivity dysfunction over long-term microelectrode implantation periods. This new knowledge could contribute to the development of targeted therapeutic strategies aimed at improving the health of the tissue surrounding brain implants and potentially inform engineering of adaptive decoders as the FBR progresses. Our study’s understanding of the dynamic changes in the functional network over time opens the door to developing interventions for improving the long-term stability and performance of intracortical microelectrodes.

## 1. Introduction

Intracortical microelectrodes enable neural signal recording and electrical microstimulation in brain-computer interface (BCI) systems. These systems have applications in both basic understading of normal brain activity as well as in clinical prostheses for restoring motor control and sensory perceptions [1-9]. For example, implanted microelectrodes can decode brain signals to drive prostheses in human patients [10], while electrical stimulation delivered by these devices helps modulate sensory perceptions in neural prosthetics [4, 11, 12]. Despite the promising role of intracortical microelectrodes, challenges remain regarding their sensitivity to record brain signals and regarding the stability of the microelectrode recordings, due to changes in network activity often attributed to ‘plasticity’ [13-20]. This challenge is represented by the difficulty in maintaining BCI decoders over time [3, 13, 14, 17, 21-24]. In addition, studies of rodents, non-human primates and human subjects have reported changes in signal quality possibly due to material and mechanical failures, such as material corrosion, degradation and/or tip breakage [25-27], or likely due to neuroinflammation and foreign body responses in surrounding tissue were reported [25, 28]. Despite knowing that implantation-related gliosis and neuronal loss contribute to the long-term decline in signal sensitivity [28, 29], there are still gaps in our understanding of how these factors specifically affect functional connectivity in brain circuits near implanted microelectrodes over time, separate from plasticity changes associated with learning.

Previous work showed that microelectrode implantation produces acute local inflammation that can affect the tissue homeostasis critical for information processing [28, 30]. The insertion of stiff, silicon-based microelectrodes disrupts the integrity of the blood-brain barrier, resulting in the infiltration of blood cells and plasma proteins [31-34]. This, in turn, recruits activated microglia and reactive astrocytes to migrate near the implanted microelectrode, leading to the formation of an encapsulating glial scar [35, 36]. The glial scar acts as a physical barrier for the exchange of ions and charged solutes between neurons and recording sites, consequently decreasing the signal-to-noise ratio thus affecting signal detectability [28, 37].

Moreover, microelectrode implantation may increase oxidative stress, pro-inflammatory cytokines, and glutamate in the local microenvironment of the electrode, affecting neuronal survival and synapse integrity [28, 38]. The consequent silencing or degeneration of nearby neurons may impair synaptic communication and neuronal activity [9, 31, 39, 40], likely contributing to diminished neural signal detection [9]. Additionally, oligodendrocytes and myelin, which are crucial for action potential conduction and to metabolically support axon function, are susceptible to injury during implantation, leading to progressive axonal degeneration and limited axonal regeneration capacity [41]. Thus, impaired oligodendrocyte and myelin function can also reduce the signals detected by the microelectrodes [42, 43].

The brain’s information processing complexity arises from its diverse anatomical structures and functional connectivity [45]. Interconnected laminar structures with distinct local networks contribute to this complexity [46]. Visual cortical circuits are thought to process information sequentially, visual information entering Layer 4 (L4), progressing to Layers 2/3 (L2/3) for integration and complex processing, and finally reaching Layer 5/6 for its output [46] to subcortical structures like the hippocampus via white matter tracts [47]. Throughout these layers, a delicate balance between excitation and inhibition is vital for circuit function. Excitation/inhibition imbalance disrupts network stability leading to behavioral impairments [48]. Interestingly, microelectrode implantation injury increases expression of excitatory neurotransmission markers within the initial days, shifting towards increased expression of inhibitory neurotransmission markers after four weeks [49, 50]. These changes suggests that the circuit function near the microelectrode is altered during long-term implantation, possibly explaining, in part, why decoders need to be frequently updated [13, 14]. Exploring the evolution of functional connectivity within and across laminar structures after implantation may generate crucial information for understanding the chronic decline in performance of implanted microelectrode devices.

Here, we aimed to investigate the changes in functional network connectivity in the cortex and hippocampal CA1 during long-term microelectrode implantation. We used multisite Michigan-style microelectrodes spanning cortical layers and the hippocampal CA1 area to record spontaneous activity and activity evoked by a visual stimulus. Our extracellular recordings detected action potentials and local field potentials (LFP) which reflect summed synaptic activity near the microelectrode [51-53]. We used cross-frequency LFP synchronization [54, 55], and action potential entrainment to LFP oscillatory activity [56], to quantify functional network connectivity. Our findings revealed depth-dependent changes in intralaminar network connectivity. Superficial Layers 2/3, rather than Layer 5, exhibited a rapid loss of detected activity for putative inhibitory neurons at 2 weeks post-implantation, whereas activity of putative excitatory neurons showed a continuous decline in firing rate without a loss of neuron detection. Furthermore, the impairment of cortical interlaminar connectivity near the microelectrode was direction-dependent. During chronic implantation, excitatory connectivity decreased in the L2/3-to-L5 descending direction but was heightened in the reverse L5-to-L2/3 direction. These findings provide the first evidence for changes in functional network connectivity within and across laminar structures during long-term microelectrode implantation. Characterizing these changes may help elucidating the mechanisms underlying the decline in activity detected by chronically implanted microelectrodes and may facilitate the strategic development of adaptive BCI decoders for longitudinal tissue reactions.

## 2. Methods

### 2.1. Intracortical microelectrode implantation surgery

C57BL6 mice (n=8, 4 males and 4 females, 11-13 weeks old, Jackson Laboratory, Bar Harbor, ME) were used in this study. A single shank 100 μm site spacing Michigan-style microelectrode (A16-3 mm-100-703-CM15, NeuroNexus, Ann Arbor, MI) was implanted in the left primary visual cortex of each animal. Surgical procedures were described in detail in previous publications [42]. Animals were anesthetized by intraperitoneally (I.P.) administration of xylazine (7 mg/kg)/ketamine (75 mg/kg) mixture. After fixing the animal onto the stereotaxic frame, hair, skin, and connective tissue were removed above the implantation site. Vetbond was applied to dry the skull for dental cement adhesion. Three stainless steel bone screws (Fine Science Tools, British Columbia, Canada) were inserted, one over each of the motor cortices and one over the contralateral visual cortex, and then secured with a layer of dental cement. The electrode ground was wired to the bone screw over the ipsilateral motor cortex and the reference was wired to the bone screws over the contralateral motor and visual cortex. A 1mm square craniotomy centered at 1.8 mm lateral to midline and 2.7 posterior to bregma was drilled. During the procedure, saline was periodically applied to prevent any thermal damage. Once the craniotomy was opened, a microelectrode was perpendicularly inserted at 15 mm/s until the tip of the microelectrode shank was sitting at 1600 um below the surface. Subtle depth adjustments were manually made to confirm the last electrode site was inserted into the brain. The craniotomy was filled with sealant Kwik-Sil and then fully covered by dental cement to seal the headcap. Body temperature and oxygen flow were maintained over the entire surgery. The animal received post-operation analgesic treatment with ketofen (I.P., 5 mg/Kg) on the surgery day and the following two consecutive days. All experimental procedures were conducted following approval by the University of Pittsburgh, Division of Laboratory Animal Resources, and Institutional Animal Care and Use Committee in accordance with the standards for humane animal care as set by the Animal Welfare Act and the National Institutes of Health Guide for the Care and Use of Laboratory Animals.

### 2.2. Electrochemical impedance spectroscopy

Electrochemical impedance spectroscopy of the implanted microelectrodes was conducted within a grounded Faraday cage. The impedance of the implanted microelectrodes was measured and verified to ensure device functionality prior to electrophysiological recording sessions. Animals were awake and head-fixed to a rotating platform while an Autolab potentiostat with 16-channel multiplexer (PGSTAT 128N, Metrohm, Netherlands) was connected to the animal head-stage. Impedance spectra were obtained for each channel using a 10 mV sine wave spanning from 10 Hz to 32 kHz. The 1 kHz impedance value was reported for each animal for each day.

### 2.3. Electrophysiological recording

Animals were awake and head-fixed in a grounded Faraday cage to prevent environmental noise during electrophysiological recordings. Spontaneous recordings were conducted with animals sitting on a rotating platform in a dark environment. Visually evoked activity was measured with animals facing a monitor (V243H, Acer. Xizhi, New Taipei City, Taiwan) to span a visual field of 60° wide and 60° high in the contralateral eye . The visual stimulation paradigm consisted of a drifting gradient of black and while solid bars (MATLAB Psychophysics toolbox) and was synchronized with the recording system (RZ2/PZ5, Tucker–Davis Technologies, Alachua FL) at 24,414 Hz. Details of this visual stimulation paradigm are described in [42, 57].

### 2.4. Electrophysiology data analysis

#### 2.4.1. Depth alignment

Current source density (CSD) plots were used to identify the laminar depth along the implanted microelectrode shank. A 2nd order Butterworth filter at 0.4-300 Hz was applied to extract the LFP data stream. The CSD heatmap was plotted by computing the average evoked (stimulus-locked) LFP for each electrode site, smoothing the signal across all electrode sites (1-dimensional line fit), and then calculating the second spatial derivative. The CSD revealed a negative LFP polarity (inward current sink) within 100 ms after visual stimuli onset, which corresponded to the L4 cortical input. The cortical alignment corresponding to CSD-defined L4 was performed in each animal for each day.

#### 2.4.2. Single-unit (SU) Classification

The raw data was passed through a Butterworth filter from 0.3-5 kHz to produce spiking data. A threshold of 3.5 standard deviations below the mean was applied to isolate potential SU activity. Then SU waveforms were manually sorted based on the quality and shape of neuronal waveforms, auto-correlograms, and peri-stimulus time histograms (PSTH) with 50 ms bins as previously described [42, 57]. The sorted SU waveforms were classified according to the spike width, defined as the latency between the trough and peak of the SU waveform [58]. The putative excitatory neurons were experimentally defined as the SU waveforms with spike width ≥ 0.41 ms, and putative inhibitory neurons were defined as the SU waveforms with spike width < 0.41 ms.

#### 2.4.3. Cross frequency LFP synchronization: phase amplitude coupling (PAC)

The functional network connectivity in the brain circuit was quantitatively described by the LFP oscillatory synchronization across different frequencies using the phase-amplitude coupling (PAC) measurements [54]. PAC defined the degree of LFP cross-frequency synchronization for how well the phase of low frequency oscillations (4-7.5 Hz) drives the amplitude of high frequency oscillations (30-90 Hz), which is calculated by the modulation index (MI).

The PAC modulation index was calculated based on Kullback–Leibler (KL) formula [54, 59]. First, raw signal was bandpass filtered to specific LFP frequencies, . The Hilbert transform was used to obtain the time series of the phase component from the slow frequency LFP activity in Channel *X*, which was denoted as *Φ*_*X*_ (*t, f*_*X*_)(*t* = time, *f*_*X*_ = phase frequencies, 4, 4.5, 5 … 7.5 Hz). The time series of the amplitude component of the high frequency LFP activity in Channel *Y* was extracted by Morlet wavelet transform (5 cycles) for each frequency (*f*_*Y*_ = amplitude frequencies, 30, 30.5, 31… 90 Hz), which was denoted as *A*_*Y*_ (*t, f*_*X*_).

Next, the composite time series was constructed *Φ*_*X*_ (*t, f*_*X*_), *A*_*Y*_ (*t, f*_*Y*_) so that the amplitude of high frequency LFP oscillation in Channel *Y* was given at each phase of slow frequency LFP activity in Channel *X*. Then the phases *Φ*_*X*_ (*t, f*_*X*_) were binned every 18° (*N* = total number of phase bins, 20), and the amplitude *A*_*Y*_ (*t, f*_*X*_) at each phase bin (*i*) was averaged as 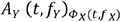 (*i*).Finally, the amplitude 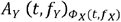, was normalized by the sum over all bins

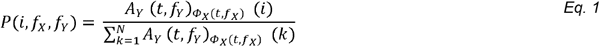

where *P*(*i*,*f*_*X*,_ *f*_*Y*_) represented the normalized amplitude distribution over phases. The normalized amplitude distribution *P*(*i*,*f*_*X*,_ *f*_*Y*_) would be uniform if there was no PAC between Channel *X* and *Y*. Thus, the level of deviation of from the uniform distribution indicated the existence of PAC, which was measured by joint entropy (*H*(*f*_*X*,_ *f*_*Y*_)).

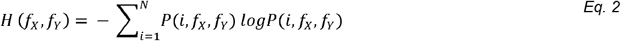

The joint entropy reaches its maximum (*H* o =*log N*^2^) when *P*(*i*,*f*_*X*,_ *f*_*Y*_)is uniform. Then, KL distance formula was applied to calculate the difference between *H*(*f*_*X*,_ *f*_*Y*_)and *H* o. Thus, the MI of PAC was calculated by dividing the KL distance from the uniform distribution *H* o.

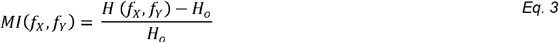

The range of MI is between 0 and 1. When Channel *X* = Channel *Y*, a larger MI value of PAC showed a stronger LFP synchronization coupling between the low frequency phase and the high frequency amplitude within the same layer, suggesting a higher activation level of the intralaminar network. When Channel *X* ≠ Channel *Y*, a larger MI value of PAC demonstrated an increased LFP synchronization coupling between the low frequency phase in the upstream layer and the high frequency amplitude in the downstream layer, reflecting an elevation in directional interlaminar connectivity.

#### 2.4.4. Spike Entrainment to LFP oscillation

As an alternative measurement for functional network connectivity, SU entrainment to LFP oscillation quantifies a directional coupling across different brain depths. Spike entrainment to LFP oscillation quantitatively describes a relationship between the timing of SU spikes and the phases of LFP [59]. The LFP oscillations are presumed to precisely organize the neuronal firing at a particular phase during communication between brain structures [56]. In this way, LFP oscillation has been considered as the oscillatory synchronized input to nearby neurons.

The modulation index (MI) of spike entrainment to LFP oscillation reflected the level of synchronization coupling between spike activity of the sorted SU and LFP oscillation at a specific frequency. The details of SU entrainment to LFP oscillation has been described previously [59]. First, raw signal of Channel *A* was bandpass filtered to specific LFP frequencies, (see below for ranges). Then, a Hilbert transform was performed to extract the time series of the phase component from the LFP activity, which was denoted as *Φ*_*X*_ (*t, f*_*A*_), where *t* represented the recording time and *f*_*A*_ represented phase frequencies (4, 4.5, 5 … 90 Hz). Then, the raw signal of Channel *B* was filtered from 0.3-5 kHz and a threshold of 3.5 standard deviations below the mean was applied to identify spike activity. The time series of spike activity was denoted as *S*_*B*_ (*t*). If there is a spike at *t*_*i*_, then *S*_*B*_ (*t*_*i*_) = 1. Otherwise, *S*_*B*_ (*t*_*i*_) = 0.

Next, the composite time series (*Φ*_*A*_ (*t, f*_*A*_), *S*_*B*_ (*t*)) was constructed to give the occurrence of spike activity in Channel *A* at phases of Channel *B* LFP oscillation. Then, the phases *Φ*_*A*_ (*t, f*_*A*_) were binned every 18° (*N* = total number of phase bins, 20), and the number of spikes *S*_*B*_ (*t*) over each phase bin (*j*) was averaged (denoted as # *spike* (*j*)). Finally, the probability distribution of spike activity over phases for each frequency *f*_*A*_ was calculated as

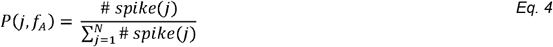

Next, the joint entropy *H*(*f*_*A*_) was calculated as

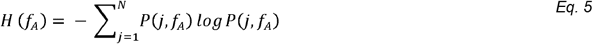

Similar to PAC, the MI for spikes in channel *B* entraining to LFP oscillation in channel *A* was calculated using the KL equation:

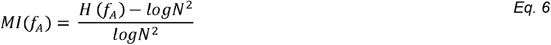

The *MI*(*f*_*A*_) of spike entrainment to LFP for each SU was normalized to its maximum value across the frequency *f*_*A*_. Then, a heatmap was constructed by the normalized MI of all sorted SU waveforms across frequencies to identify the LFP frequency range (*f*_*k*_= *f*_1_, *f*_2_,… *f*_*n*_) where the most SU waveforms were strongly entrained. If Channel *A* = Channel *B*, the MI measured the level of spike entrainment to the intralaminar LFP oscillation. If Channel *A* ≠ Channel *B*, the MI of spike entrainment to the interlaminar LFP oscillation reflects the directional network connectivity across cortical depth.

Once the LFP frequencies (*f*_*n*1_ − *f*_*n*2_) were identified, the raw data in Channel *A* was bandpass filtered to *f*_*n*1_ − *f*_*n*2_ frequencies. Then a Hilbert transform was applied to extract phase angles at each spike activity at Channel *B*. The directional statistics were performed using MATLAB Circular Statistics Toolbox. The mean resultant vector was measured, which reflected the mean phase angle of spike entrainment for that SU. The resultant vector length was reported to describe a spread of the individual phase angles about the mean angle. The sorted SU was considered entrained if the resultant vector length was significantly deviated from the uniformity threshold set by Rayleigh test (*p <* 0.05) [60]. Only the significantly entrained SU were statistically compared between different time periods of implantation.

### 2.5. Immunohistochemistry

16 weeks after microelectrode implantation, mice were perfused transcardially with 1x PBS and then 4% paraformaldehyde (PFA). The brain tissues were then postfixed with 4% PFA overnight at 4°C, immersed in 30% sucrose for rehydration, and finally sectioned coronally into 25 μm slices until the probe trace was visualized. Standard immunohistochemical staining was performed. Following heat-induced antigen retrieval (0.1 M citric Acid, 0.1 M sodium citrate) and endogenous peroxidase blocking, the tissue was covered by 0.1% Triton-X with 10% normal goat serum in PBS at room temperature for 1 h. Primary antibodies GAD67 (1:500, Abcam, ab213508)and CamKiiα (1:100, Abcam, ab22609) were incubated overnight at 4°C. Then the secondary antibodies (Nissl 435/455, Thermo Fisher, N-21479, donkey anti-rabbit 488, Abcam, ab150061, donkey anti-mouse 568, Abcam, ab175700, donkey anti-goat 647, Abcam, ab150135) in concentration of 1:500 were applied. After incubation, slides were washed with PBS and mounted with cover glass over Fluoromount-G media (SouthernBiotech, #0100-20). The 16-bit, 1024 × 1024 pixels (635.9 × 635.9 μm), 20x TIFF images were captured using a confocal microscope (FluoView 1000, Olympus, Inc., Tokyo, Japan) system with an oil-immersive objective lens.

### 2.6. Statistics

Significant differences between visual evoked and spontaneous conditions were assessed using a linear mixed effect model to account for repeated measures. The model applied a restricted cubic spline with 4 knots at the 5th, 35th, 65th, and 95th percentiles of the data for nonlinear fit. In this model, the time and the condition-by-time interaction are fixed effects. The group-wise difference was significant if the 95% confidence intervals (1.96 times the standard error of the model fits) were not overlapping based on a likelihood test. The significant changes in metrics over time were assessed using a one-way ANOVA with Tukey post hoc tests. The Repeated Measures ANOVA was applied to detect significance in angular measurements across different implantation stages, using the Watson-Williams test for the resultant mean entrainment angles and the Kuiper test for the distribution of entrainment angles.

## 3. Results

### 3.1. Layer-specific characteristics of classified SU firing activity

Given the distinct patterns of anatomical connectivity across cortical layers and hippocampus, we first examined the firing activity of putative neuronal subtypes at various depths. Sixteen-channel microelectrodes were perpendicularly inserted into the brain through the primary visual cortex and to a depth of 1.6 mm so that the the tip reached hippocampus CA1 (Fig. 1A). No material failure or mechanical failure was observed in these implanted microelectrodes, and the electrochemical properties of the implanted microelectrodes remained stable over time as indicated by the non-significant differences in channel-averaged impedance (Fig. 1B, One-way ANOVA, *p* = 0.569). The recorded SU spikes are widely acknowledged to represent action potentials generated by individual neurons [61, 62]. In this study, we further classified these SU waveforms into putative excitatory or inhibitory neurons to gain detailed insights into firing activity within neuronal networks at different depths using trough-to-peak (TP) latency of all SU waveforms [58, 59, 63]. A bimodal distribution was observed with narrow waveforms peaking at 0.25 ms, consistent with previous observations for inhibitory neurons [37,38] and the wide waveforms peaking at 0.54 ms, corresponding to excitatory neurons [64]. Using a threshold of 0.41 ms, we identified a total of 440 waveforms as putative inhibitory neurons and 1,476 waveforms as putative excitatory neurons in awake mice over the entire 16-week implantation period (Fig. 1C).

**Figure 1.**
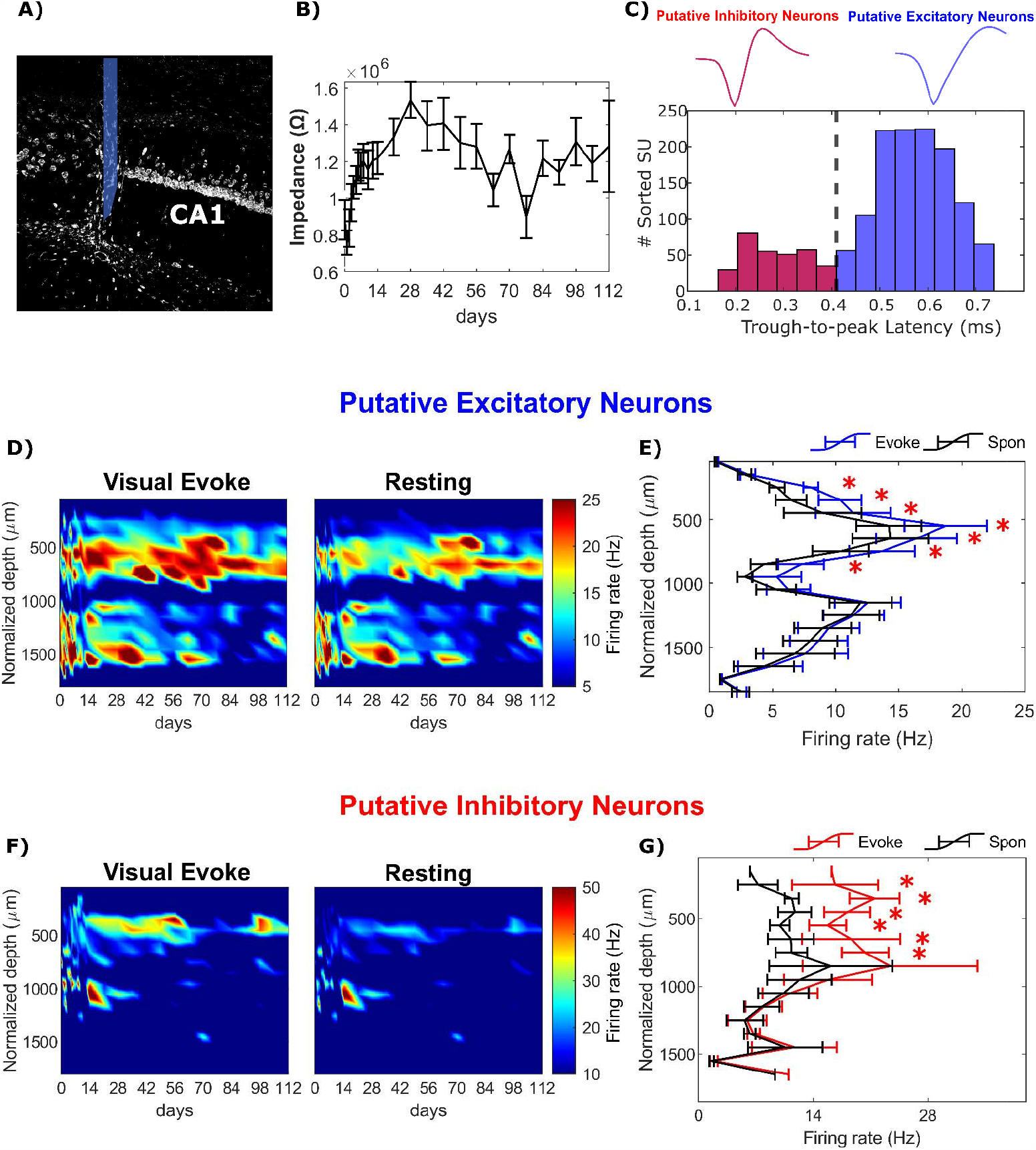
Visual stimulation activates distinct neuronal subtypes in depth-dependent manner. A) Representative Nissl+ neuronal soma staining of the hippocampal CA1 reveals the probe trace with the tip positioned below CA1. The location of the probe is labeled in blue. B) Impedance at 1 kHz was averaged across channels and compared between time points (*p* = 0.569). C) Top: Example of a representative narrow waveform classified as a putative inhibitory neuron (left) and representative wide waveform sorted as a putative excitatory neuron (right). Bottom: Single-units (SUs) were classified based on trough-to-peak (TP) latency, which followed a bimodal distribution. SU waveforms with TP latencies less than 0.41 ms were classified as putative inhibitory neurons (red), while SU waveforms with TP latencies greater than 0.41 ms were classified as putative excitatory neurons (blue). D) SU firing rate heatmaps of putative excitatory neurons during visual evoked responses (left) and resting state (right) as a function of time and depth. E) The depth profile of putative excitatory firing rate showed a significant increase in firing rate in the cortex during visual stimulation. F) SU firing rate heatmaps of putative inhibitory neurons during visual evoked responses (left) and resting state (right) as a function of time and depth. G) Putative inhibitory neurons exhibited a significant increase in firing rate in the cortex (peaked in L2/3) during visual stimulation compared to the resting state. * indicates non-overlapping 95% confidence intervals between visual evoked responses and resting state at each time point, determined using a linear mixed-effects model with likelihood ratio test..

The firing rate of putative excitatory neurons with wide spike waveform exhibited two distinct peaks along the depth axis (Fig. 1D). The first peak, with a firing rate of 18.72 ± 3.26 Hz, was observed at ∼600 μm depth corresponding to L5 neurons. This finding is consistent with previous observations of high firing rates of large L5 pyramidal neurons in cortex [65] (Fig. 1E). The second peak, with a firing rate of 12.54 ± 2.62 Hz, was observed at ∼1,200 μm depth, likely corresponds to pyramidal neurons in CA1 Stratum Pyramidale (SP) characterized by dense cell bodies (Fig. 1D-E). This firing rate was relatively higher than the firing rate of CA1 neurons as reported [66], which is likely due to multiple neurons being classified under the same waveform in extracellular recording conditions. During visual stimulation, the firing rate of putative excitatory neurons in visual cortex significantly increased compared to resting state within the depth range of 250-850 μm (*p* = 0.078, likelihood ratio test with non-overlapping 95% confidence intervals). Notably, the largest increase, approximately 31%, was observed in Layer 4 (L4), which is the primary target of excitatory thalamic afferents conveying information from the visual stimuli [67, 68]. In contrast, in the hippocampus, the visually evoked increase in the firing rate of putative excitatory neurons was limited to only around 5%, showing that visual stimulation with a drifting-bar gradient had a relatively weaker effect in activating the hippocampus CA1 region near the implanted microelectrode. Taken together, the firing rate of putative excitatory neurons displayed a significant increase in L4 during visual stimulation, but limited effects in the hippocampus CA1 region near the microelectrode.

In contrast to putative excitatory neurons, putative inhibitory neurons with narrow spike waveform from visual cortex displayed consistent firing rate across cortical depth and exhibited a higher overall firing rate. (Fig. 1F; mean firing rate of putative excitatory neurons: 11.79 Hz, mean firing rate of putative inhibitory neurons: 18.55 Hz). The increase in putative inhibitory firing rate during visual stimulation was statistically significant at depths ranging from 150-750 μm below the surface (*p* = 7.27 * 10^(-5)). However, in the hippocampus, the increase was only 3% compared to resting state (Fig. 1G). The peak increase in putative inhibitory firing rate, approximately 89% higher than resting state, occurred in L2/3. Given that GABAergic interneurons regulate signal flow and shape network dynamics [69, 70], the substantial elevation in putative inhibitory firing rate in L2/3 indicates is consistent with the critical contribution of interneuron-mediated processes in this layer, such as lateral inhibition, in cortical information processing. In summary, the putative excitatory and inhibitory neuron subtypes in visual cortex exhibited different depth profiles of firing rate during visual stimulation. Whereas the input layer L4 experienced a prominent increase in excitatory neuron activity, L2/3 showed the strongest inhibitory network activation. Interestingly, the hippocampal CA1 region did not show a significant increase in firing activity for either putative neuronal subtypes during passive drifting-bar gradient visual stimulation.

### 3.2. Patterns of Change in Cortical Laminar Networks near the Chronically Implanted Microelectrode

The depth profile of the firing rate of putative neuronal subtypes provides insights into the interconnectedness of cortical laminar structures, which exhibit distinct network functionality for information processing. Our objective was to gain better understanding of the changes in functional network connectivity in cortex over long-term microelectrode implantation. To achieve this we first examined specific cortical intralaminar networks involved in visual processing. To evaluate the impact of long-term microelectrode implantation on functionality of intralaminar networks, we examined the LFP cross-frequency synchronization, single-unit (SU) firing rates of putative neuron subtypes, and spike entrainment to intralaminar LFP. SU spiking activity specifically requires the involvement of neurons in close proximity to the electrode site [71-73] (usually within 80-140 μm). Studying LFP cross-frequency synchronization allows us to gauge the level of network activation over a broader distance, while changes in spiking rate can provide insights into the function of local network surrounding to the implanted microelectrode within about 100 μm. Since LFP oscillations are understood to reflect synchronized input to nearby populations of neurons [56, 74], analyzing spike entrainment of individual neurons to the population’s intralaminar LFP oscillation informs us about the responsiveness of the individual neurons to the surrounding network. By employing these three measurements, we aim to uncover a novel perspective on potential changes in network functionality near the chronically implanted microelectrode.

#### 3.2.1. Stable Excitatory Connectivity in L4

We started our investigation in L4 as it is the primary input-receiving layer in cortex. To investigate the functional connectivity within the intralaminar network of L4 during visual processing, we examined the LFP cross-frequency synchronization between theta (4-7.5 Hz) and gamma (30-90 Hz) oscillations within depths corresponding to L4. This theta-gamma coupling has been recently identified as a critical neural coding mechanism in cognition [75], where the phase of theta oscillation is strongly coupled with the gamma power during network activation [76]. To quantitatively assess the phase-amplitude coupling (PAC) between theta and gamma oscillatory activity, we calculated the Modulation Index (MI) for LFP synchronization. The MI measures the strength of synchronization between the low-frequency phase and high-frequency amplitude, with a range between 1, indicating strong LFP synchronization, and 0, indicating loss of synchronization. Our results revealed a robust PAC relationship between the phase around ∼4 Hz and the amplitude in the 80-90 Hz range during visual evoked recordings (Fig. 2A). Furthermore, we observed that the MI value during visual stimulation consistently remained significantly elevated compared to the MI during resting state throughout the entire 16-week chronic implantation period (Fig. 2B, *p* = 1.45 * 10 ^(-7)). Aditionally, there were no significant differences in MI values during visual stimulation across different time points (One-way ANOVA, *p* = 0.361). Our LFP findings within L4 indicate that the intralaminar network in this layer is consistently activated near the chronically implanted microelectrode in response to visual stimuli.

**Figure 2.**
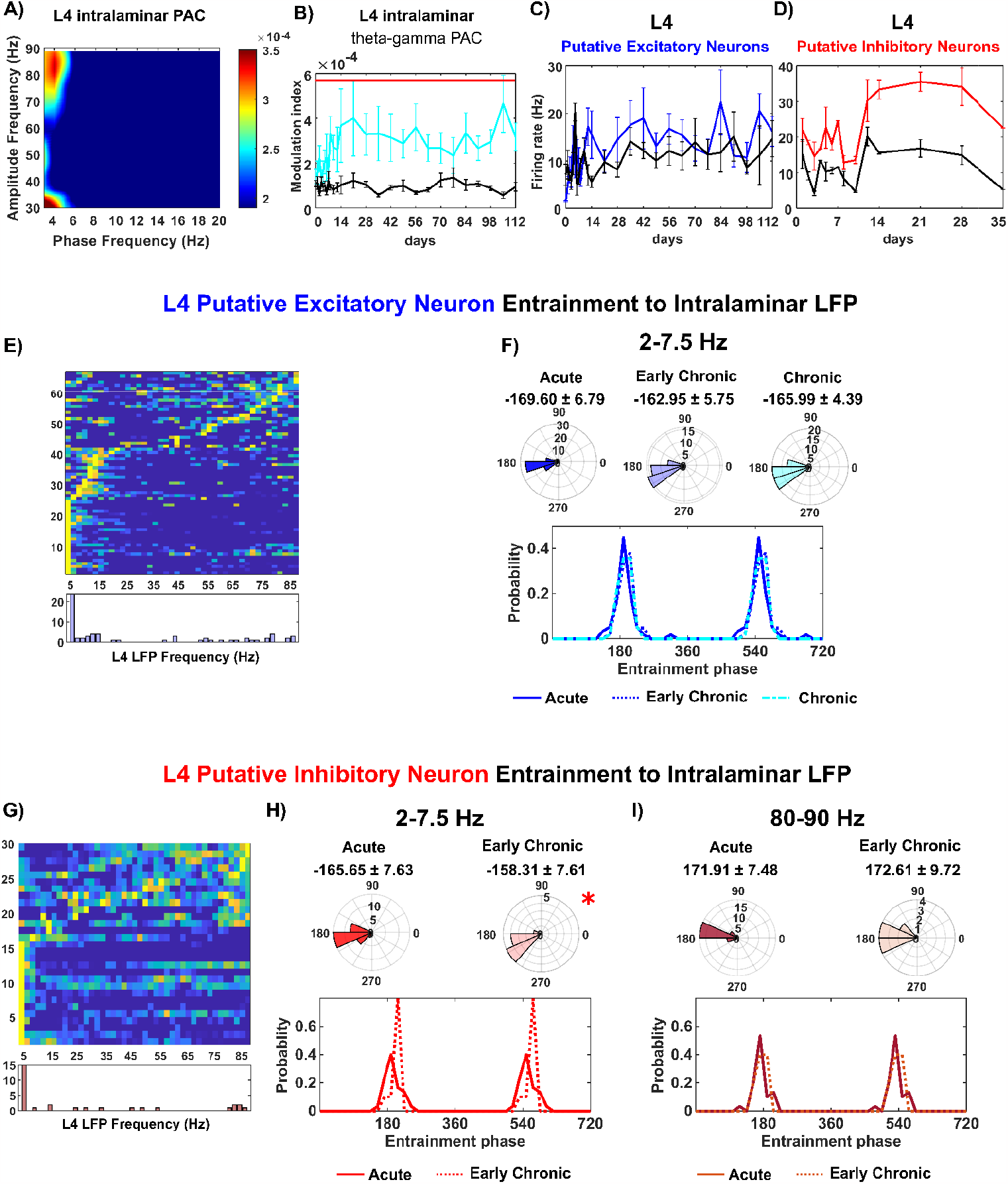
L4 maintained a stable excitatory connectivity over time but experienced loss of putative inhibitory neuron activity 5-weeks post implantation. A). Comodulogram displays the Modulation Index (MI) of phase frequency (2-20 Hz) with the amplitude frequency (30-90 Hz) in L4. Note: Strong coupling is observed between ∼4 Hz phase and the ∼80-90 Hz amplitude during visual stimulation. B). The MI index of the 4 Hz t eta phase coupling with 80-90 Hz gamma amplitude is significantly higher during visual stimulation (cyan) compared to the resting state (black) throughout the entire 16-week microelectrode implantation period. The red bar indicates significance of this difference across all time points. C). The firing rate of putative excitatory neurons during visual evoked activity (blue) and resting state (black) is plotted over time. D). The putative inhibitory firing rate is significantly higher (*p <* 0.05) during visual stimulation (red) compared to resting state (black) but detection of inhibitory neurons is lost after week 5 post-implantation. E). L4 putative excitatory neurons exhibit entrainment to intralaminar slow oscillation. The top panel shows the normalized spike-LFP entrainment MI across frequencies for each L4 putative excitatory neuron with a Rayleigh test *p*-value < 0.05. The bottom panel displays a histogram indicating that a group of putative excitatory neurons are predominantly modulated by the L4 intralaminar slow oscillation (2-7.5 Hz). F). The entrainment phase angles of L4 putative excitatory neurons to intralaminar slow 2-7.5 Hz oscillation are shown in a polar histogram (top) and line plot (bottom). The mean phase angle of entrained L4 excitatory neurons is reported for the acute (0-2 weeks), early chronic (3-8 weeks), and chronic (9-16 weeks) implantation periods. G) The top panel demonstrates the normalized spike-LFP entrainment MI value for each significantly entrained putative inhibitory neuron (Rayleigh test p-value < 0.05) across frequencies. The bottom panel shows a histogram indicating that putative inhibitory neurons are primarily entrained to two frequencies: the slow 2-7.5 Hz oscillation and the high 80-90 Hz frequency. The polar histograms and line plots illustrate the entrainment phase of L4 putative inhibitory neurons to the slow 2-7.5 Hz frequencies (H) and the 80-90 Hz high frequencies (I) of L4 intralaminar LFP at the acute (0-2 weeks) and early chronic (3-8 weeks) implantation stages. Spike-LFP entrainment analysis was not performed during the chronic (9-16 weeks) stage due to the loss of L4 putative inhibitory neuron detection. The red bar indicates non-overlapping 95% confidence intervals between visual evoked responses and resting state at each time point, as determined by a linear mixed effects model with likelihood ratio test. The asterisk (*) indicates significant differences between the phase angular distribution in terms of resultant mean angles (Watson-Williams test), distribution variability (Kuiper test), or both.

Then, we examined the firing rate of individual L4 SU waveforms near the microelectrode. The firing rate of putative excitatory neurons in L4 showed a noticeable increase during visual stimulation compared to the resting state, indicating their involvement in L4 activation (Fig. 2C). Moreover, the firing rate of putative excitatory neurons in L4 remained stable over time (Two-way ANOVA with Tukey post hoc, *p* = 0.307). In the case of putative inhibitory neurons in L4, visual stimulation led to a significant elevation in their firing rate compared to the resting state (Fig. 2D, *p =* 0.0034), emphasizing the crucial role of inhibitory neurons in L4 activation. However, starting from week 5 post-implantation, there was a decline in the detection of sufficient putative inhibitory neurons in L4. It is possible that local interneurons detected as the putative inhibitory neurons here became silent or injured due to microelectrode implantation. However, those interneurons in distal regions that are outside the detection range of the microelectrode are likely still functioning and generate the theta and gamma rhythms (Fig. 2A-B).

Next, we investigated whether the activity of individual neurons during the intralaminar LFP oscillatory input was influenced by the chronic implantation injury. While the overall firing rate of L4 putative excitatory neurons remained stable over time, it was important to determine if these neurons were able to effictively synchronize (phase lock) their activity with LFP oscillations. We first focused on the entrainment of L4 putative excitatory neurons to slow 2-7.5 Hz LFP oscillations. Remarkably, a substantial portion (nearly 42%) of L4 putative excitatory neurons exhibited a significant entrainment to this slow 2-7.5 Hz LFP oscillation (Rayleigh test, *p <* 0.05; Fig. 2E). These neurons consistently aligned their spiking activity with the bottom of the slow oscillation, and their distribution variability of entrainment phases and mean preferred phase did not exhibit significant differences over time (Fig. 2F; Watson–William’s test, *p >* 0.05; Kuiper test, *p >* 0.05). Thus, the spike timing of L4 excitatory neurons consistently synchronized with the trough of the slow oscillation throughout the entire chronic implantation period, suggesting stable activity during LFP oscillations of L4 excitatory neurons.

We further investigated the activity of L4 putative inhibitory neurons by examining their spike entrainment to intralaminar LFP. Our analysis revealed that 53.3% of L4 putative inhibitory neurons exhibited significant entrainment to the intralaminar 2-7.5 Hz oscillation, and 20% showed significant entrainment to 80-90 Hz gamma oscillation (Fig. 2G, Rayleigh test, *p <* 0.05). However, unlike putative excitatory neurons, the L4 putative inhibitory neurons displayed significant changes in distribution variability of entrainment phases of slow 2-7.5 Hz oscillation during the early chronic stages (3-8 wks) compared to the first two weeks (Fig. 2H; Kuiper test, *p <* 0.05). This significant change in spike-LFP coupling of L4 putative inhibitory neurons suggests an early deficit in the ability of L4 inhibitory neurons to respond to intralaminar slow oscillations. On the other hand, L4 putative inhibitory neurons exhibited no significant differences in mean phase or phase distribution variability of 80-90 Hz gamma oscillation across different implantation stage (Fig. 2I; Watson–William’s test, *p >* 0.05; Kuiper test, *p >* 0.05). The firing activity of L4 putative excitatory neurons and their relationship with oscillatory LFP input remained preserved over time, indicating the robustness of the L4 intralaminar excitatory network during the chronic 16-week implantation period. However, the putative inhibitory neurons in L4 exhibited an early deficit in spike entrainment to slow oscillation, suggesting L4 inhibitory neurons experienced an impairment in synchronization of firing activity to surrounding oscillation. This dysregulation was followed by a detection loss of L4 putative inhibitory neurons, the temporal sequence indicating an early dysfunctionality of these neurons near the chronically-implanted microelectrode. Taken together, these results indicate that the primary input-receiving L4 maintained the excitatory functional connectivity yet experienced an intralaminar inhibitiory activity.

#### 3.2.2. Rapid Decline in L2/3 Network Activity

In the visual cortex, it is well-acceptedthat after most of the sensory information enters L4, it is then transmitted to L2/3, and finally to L5/6 [77]. Given the gradual loss in the detection of local inhibitory neuronal activity in L4, we then asked whether, when, and how the activity of the downstream L2/3 intralaminar network changes during the 16-week microelectrode implantation period. First, we performed an analysis of LFP cross-frequency synchronization, which was focused on the coupling between theta phase and gamma amplitude in L2/3 LFP oscillations. The MI of the L2/3 theta-gamma PAC demonstrated a strong coupling between ∼5 Hz theta phase and ∼65-90 Hz gamma amplitude during visual stimulation (Fig. 3A). In comparing the visually-evoked responses with the resting state, we observed significant differences in MI values. This underlines that visual stimulation results an increased LFP synchronization in L2/3 and thus indicates an enhanced intralaminar network connectivity relative to resting state. Interestingly, we observed a gradual decline in the MI value of L2/3 theta-gamma PAC during visual stimulation, reaching a comparable level to the resting state at week 8. However, starting from week 13 post-implantation, the MI values significantly increased again (Fig. 3B, *p* = 1.99 *10^(-12)). The initial decline in MI during visual stimulation suggests a functional impairment in the L2/3 intralaminar network near the microelectrode, while subsequent increase in MI values suggests a chronic enhancement. This longitudinal changes in L2/3 intralaminar connectivitypotentially indicates a remodeling process of functional network connectivity. These findings shed light on the dynamic changes in the L2/3 intralaminar network near the chronically implanted microelectrode. The impairment followed by potential remodeling suggests a complex interplay between the microelectrode and the functional network connectivity in L2/3.

**Figure 3.**
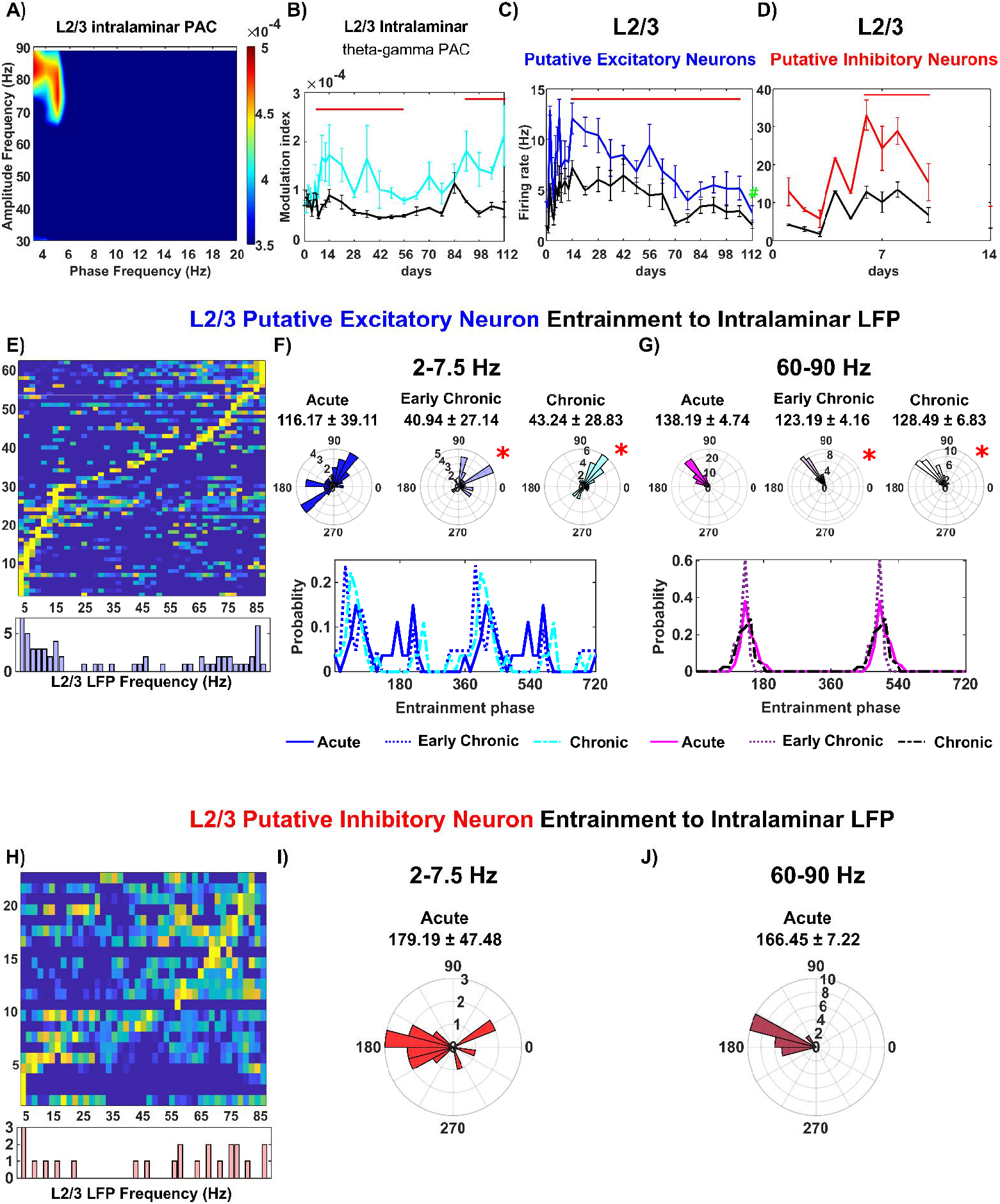
Electrode implantation leads to a loss of L2/3 putative inhibitory neurons and a significant reduction in the firing rate of putative excitatory neurons. A) Comodulogram of L2/3 intralaminar theta-gamma modulation index (MI) showing the phase frequency (2-20 Hz) with the amplitude frequency (30-90 Hz). Visual evoked responses induce a robust coupling between ∼5 Hz phase and the ∼65-90 Hz amplitude. B) Time series of L2/3 theta-gamma phase-amplitude coupling (PAC) during visual evoked responses (cyan) and resting state (black) over time. C) Putative excitatory neurons in L2/3 experience a substantial decrease in firing rate during visual evoked activation. Visual evoked responses significantly increase the firing rate of putative excitatory neurons compared to the resting state (*p <* 0.05). D) Putative inhibitory neurons in L2/3 significantly increase their firing rate during visual evoked responses compared to the resting state, but the detection of these units is lost after week 2 post-implantation. E) Putative excitatory neurons in L2/3 show entrainment to intralaminar slow and fast oscillations. The top panel shows normalized spike-LFP entrainment modulation indices (MIs) across frequencies for each putative excitatory neuron in L2/3 with a Rayleigh test p-value < 0.05. The bottom panel presents a summary histogram indicating that a cluster of putative excitatory neurons in L2/3 is mostly entrained to intralaminar slow oscillations (2-7.5 Hz), while the other cluster is mostly entrained to high-frequency oscillations (60-90 Hz). F) Distribution of phases of putative excitatory neurons in L2/3 significantly entrained to 2-20 Hz slow frequency oscillations at three different implantation stages. The top panel shows a polar histogram with the mean phase reported, while the bottom panel displays a line plot. G) Distribution of phases of putative excitatory neurons in L2/3 significantly entrained to 60-90 Hz high-frequency oscillations at different implantation stages. The top panel shows a polar histogram, and the bottom panel presents a line plot. H) Putative inhibitory neurons in L2/3 during the acute 0-2 weeks of implantation are entrained to both slow 2-7.5 Hz oscillations and 60-90 Hz oscillations, phase-locked to the trough of these slow (I) and fast (J) oscillations. The red bar indicates non-overlapping 95% confidence intervals between visual evoked responses and resting state at each time point, determined using a linear mixed-effects model with likelihood ratio test. The # symbol in 2C indicates significant differences over time determined by One-way ANOVA with Tukey post hoc tests. The * symbol indicates significant differences between phase angular distribution in either the resultant mean angles (Watson–Williams test) or distribution variability (Kuiper test), or both.

The firing activity of both L2/3 putative excitatory and inhibitory neuronal subtypes exhibited a progressive decline. Specifically, the firing rate of L2/3 putative excitatory neurons showed a substantial declined over the 16 weeks post-implantation period, with a significant reduction observed at week 16 compared to weeks 1-8 (Fig. 4C, *p <* 0.01). This impaired firing activity indicates progressive impairment of the L2/3 excitatory network. Furthermore, L2/3 putative inhibitory neurons displayed a significant increase in visually evoked firing rate compared to resting state (Fig. 3D, *p* = 7.82 * 10^(-6)). However, the detection of sufficient L2/3 putative inhibitory neurons was not possible after week 2 post-implantation, suggesting that inhibitory neurons in L2/3 were susceptible to acute implantation injury. This dysfunction of L2/3 putative excitatory and inhibitory neuronal firing activity indicates that the local L2/3 network adjacent to the microelectrode was compromised, which is further supported by the evidence of immunohistochemical staining (Fig.8).

**Figure 4.**
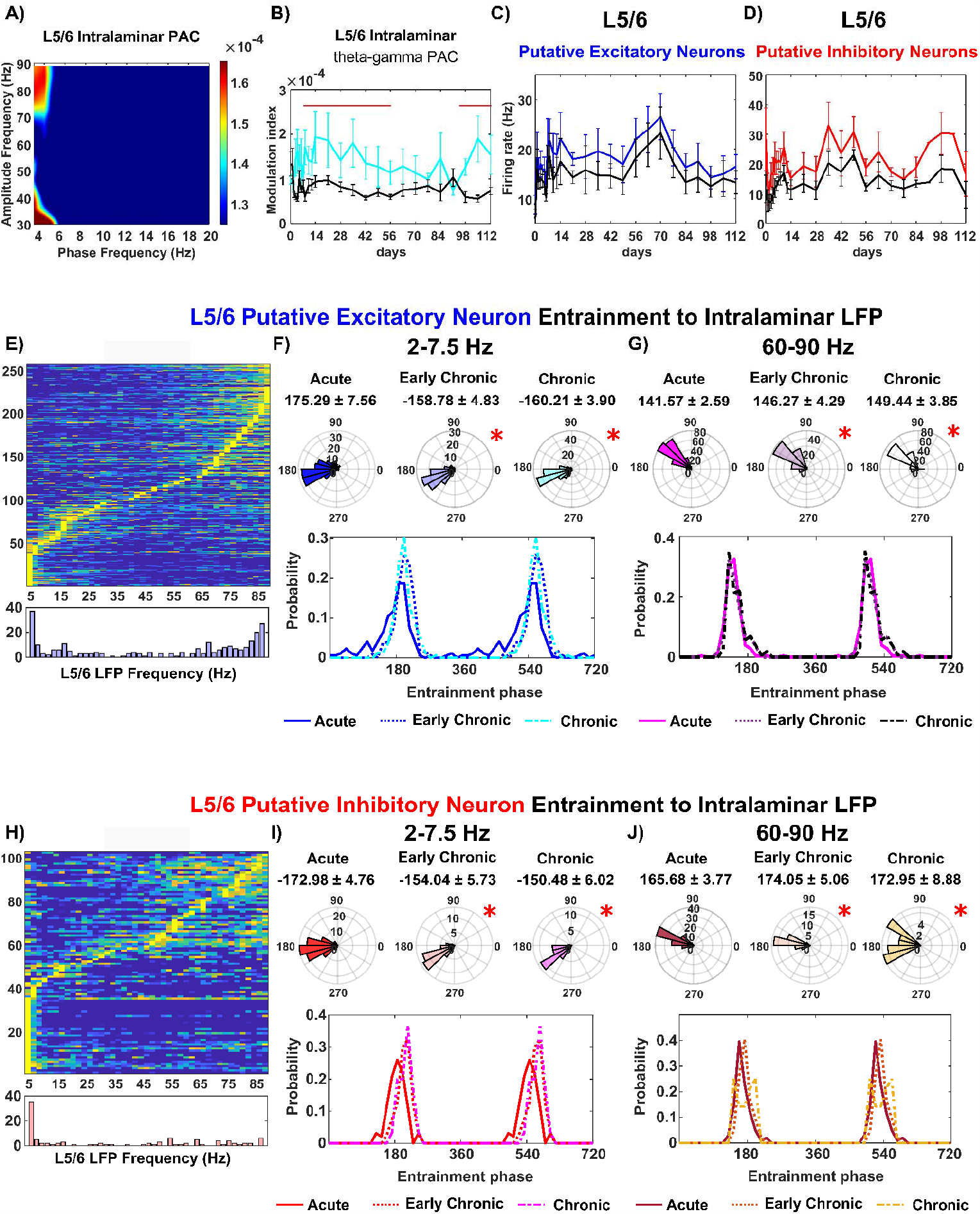
Impaired intralaminar connectivity near the chronically implanted microelectrode in cortical output layer L5/6. A) Comodulogram of L5/6 intralaminar theta-gamma modulation index (MI) showing a strong phase-amplitude coupling between ∼4 Hz phase and the ∼70-90 Hz amplitude during visual evoked responses. B) Plot demonstrating the robust modulation index (MI) between visual evoked responses (cyan) and resting state (black) over time. C) Firing rate of putative excitatory neurons in L5/6 was significantly higher during visual evoked responses compared to the resting state (linear mixed model, *p <* 0.05). D) Firing rate of putative inhibitory neurons in L5/6 was significantly higher during visual evoked responses compared to the resting state (linear mixed model, *p <* 0.05). E) Putative excitatory neurons in L5/6 were entrained to intralaminar LFP oscillations. The top panel shows normalized spike-LFP entrainment MI across frequencies for each putative excitatory neuron in L5/6 with a Rayleigh test p-value < 0.05. The bottom panel presents a summary histogram indicating that a subpopulation of putative excitatory neurons in L5/6 was entrained to intralaminar slow 2-7.5 Hz oscillations, while another subpopulation was entrained to 60-90 Hz oscillations. F) Preferred phase distribution of putative excitatory neurons in L5/6 entraining to 2-7.5 Hz oscillations over different implantation stages. The top panel shows a polar histogram, and the bottom panel presents a line plot. G) Preferred phase distribution of putative excitatory neurons in L5/6 entraining to 60-90 Hz oscillations over different implantation stages. The top panel shows a polar histogram, and the bottom panel presents a line plot. H) Putative inhibitory neurons in L5/6 were entrained to two frequencies of intralaminar LFP oscillations: one cluster of 2-7.5 Hz frequencies and another cluster of 60-90 Hz frequencies. I) Preferred phase distribution of putative inhibitory neurons in L5/6 entraining to 2-7.5 Hz oscillations over different implantation stages. J) Preferred phase distribution of putative inhibitory neurons in L5/6 entraining to 60-90 Hz oscillations over different implantation stages. The top panel shows a polar histogram, and the bottom panel presents a line plot. The red bar indicates non-overlapping 95% confidence intervals between visual evoked responses and resting state at each time point, determined using a linear mixed-effects model with likelihood ratio test. The * symbol indicates significant differences between phase angular distribution in either the resultant mean angles (Watson–Williams test) or distribution variability (Kuiper test), or both.

Furthermore, our analysis of spike entrainment to the LFP in L2/3 sheds light on the changes of neuronal communication near the microelectrode. Specifically, putative excitatory neurons in L2/3 exhibited a preference for entrainment to intralaminar 2-7.5 Hz theta frequencies (19.7%) as well as 60-90 Hz gamma frequencies (36%) (Fig. 3E). However, these neurons were unable to maintain an organized distribution of entrainment phase to either frequency band. Significant differences were detected in both the mean as well as the variability of entrainment phase angles during the early chronic phase (3-8 weeks) and chronic phase (9-16 weeks) compared to the acute phase (0-2 weeks) (Fig. 3F, Watson–William’s test *p <* 0.05; Kuiper test, *p <* 0.05). Additionally, the mean phase of L2/3 putative excitatory spike entrainment to intralaminar gamma oscillation (60-90 Hz) showed a significant shift in the mean phase since early chronic phase (3-8 weeks) (Fig. 3G, Watson–William’s test, *p <* 0.05). Visual inspection showed a shift in probability of entrained phase from ∼130 ° during the acute phase (0-2 weeks) to ∼160 ° during the chronic phase (9-16 weeks). Yet, there was no significant correlation of the phase angles with the firing rate of L2/3 putative excitatory neurons (Pearson’s coefficient: 0.166, *p* = 0.154), which suggested the loss of synchronization . Together, these findings indicate that putative excitatory neurons in layer 2/3 progressively lose the ability to fire consistently in phase with intralaminar LFP oscillations over time.

For putative inhibitory neurons, the analysis of spike-LFP entrainment to intralaminar LFP oscillations were only reported at the acute phase (0-2 weeks) due to a loss of detectable putative inhibitory neurons (Fig. 3H). We observed that the ability of L2/3 putative inhibitory neurons to entrain to multiple frequencies was rapidly lost after the second week following implantation. Specifically, these L2/3 putative inhibitory neurons exhibited phase locking to the trough of slow 2-7.5 Hz oscillation (mean phase: 179.19 ± 47.48°) as well as 60-90 Hz gamma oscillation (mean phase: 166.45 ± 7.22°). While inhibitory neurons are known to play a critical role in local network communication [78], the observed loss in L2/3 putative inhibitory neurons suggests an early onset of local network dysfunction near the microelectrode. Together, these results indicate the L2/3 network experiences early impairment in close proximity to the microelectrode and undergoes progressive degeneration, implying a disruption in information processing during brain activity.

#### 3.2.3. Impaired Neuronal Entrainment of L5/6 to Intralaminar LFP Oscillation

We then investigated the network function of L5/6, which serves as the major cortical output to other cortical areas and subcortical structures [79]. Analyzing the phase-amplitude coupling (PAC) of L5/6 intralaminar local field potential (LFP) oscillations, we found that the phase of ∼4 Hz theta oscillations strongly modulated the amplitude of 70-90 Hz gamma oscillations during visual evoked responses (Figure 4A). The significant increase in the modulation index (MI) value of L5/6 theta-gamma PAC during visual evoked responses compared to the resting state (Figure 4B, linear mixed model, *p* = 2.98 * 10^(-11)) indicates the functional activation of the L5/6 intralaminar network near the microelectrode. Interestingly, the pattern of visual evoked MI over time in L5/6 resembled that of L2/3. The significantly elevated MI during visual evoked responses dropped to the level of resting state at week 8 but significantly increased again since week 14 post implantation (Fig. 4B). This suggests that the microelectrode implantation simultaneously influenced the activity of both the upstream L2/3 network and the downstream L5/6 intralaminar network.

However, the firing rate of both putative excitatory and inhibitory neuron subtypes in L5/6 remained stable over time. Both putative excitatory neurons (Figure 4C, *p* = 0.043) and inhibitory neurons (Figure 4D, *p =* 0.0027) in L5/6 exhibited a significant increase in firing rate during visual evoked responses compared to the resting state. Additionally, there were no significant differences in the firing rate of each putative neuron subtype across different time points. It is worth noting that there was sufficient detection of putative inhibitory neurons in L5/6, likely due to the increased density of inhibitory neurons distributed in the deeper cortical layers [69, 80].

Although the overall firing rates of L5/6 putative neuron subtypes remained stable over chronic implantation, it is possible that they were unable to phase-lock to the intralaminar LFP oscillatory input. Therefore, we examined the spike entrainment of L5/6 neurons to intralaminar LFP oscillation at different implantation stages. For L5/6 putative excitatory neurons, we found that 19.2% were entraining to intralaminar 2-7.5 Hz LFP oscillations, and nearly 50% were entraining to intralaminar 60-90 Hz gamma oscillation (Fig. 4E). These putative excitatory neurons in L5/6 exhibited phase-locking to the trough of the intralaminar 2-7.5 Hz oscillation during the acute 0-2 weeks, which significantly shifted in counter-clockwise direction at early chronic 3-8 weeks and chronic 9-16 weeks (Fig. 4F, Watson–William’s test, *p <* 0.05; Kuiper test, *p <* 0.05). Over time, there was an increase in the probability of preferred phase of ∼160 degree at early chronic 3-8 weeks stage (-158.78° ± 4.83°), followed by a decrease in the probability of preferred phase of ∼200 degree at chronic 9-16 weeks (-160.21° ± 3.90°). Additionally, the L5/6 putative excitatory neurons entraining to intralaminar 60-90 Hz high frequency oscillation were phase-locked near 140° in first 2 weeks (141.57° ± 2.59°). The mean of preferred phases significantly shifted counterclock-wise (Fig. 4G, Watson–William’s test, *p <* 0.05) to 146.27° ± 4.29° in early chronic 3-8 weeks and 149.44° ± 3.85° chronic 9-16 weeks. Overall, these L5/6 putative excitatory neuron altered their activity to intralaminar LFP oscillatory input during the chronic implantation period.

Meanwhile, nearly 40% of L5/6 putative inhibitory neurons were entraining to 2-7.5 Hz oscillation, and 26% were entraining to 60-90 Hz gamma oscillation (Fig. 4H). These putative inhibitory neurons in L5/6 were preferentially phase-locked to the trough of 2-7.5 Hz LFP oscillation during the acute 0-2 weeks (-172.98° ± 4.75°). However, the mean preferred phase of this entrainment was significantly shifted counterclockwise with a reduction in the probability of preferred phase in the range of ∼150°-170° (Fig. 4I). Moreover, the variability of the preferred entrainment phase became significantly narrower (*p <* 0.05) from early chronic 3-8 weeks. Additionally, when entrained to intralaminar 60-90 Hz LFP, L5/6 putative inhibitory neurons were phase-locked at 165.68° ± 3.77° in acute 0-2 weeks. However, the distribution of the preferred phase significantly increased in variability (Fig. 4J, Kuiper test, *p <* 0.05) at early chronic 3-8 weeks (174.05° ± 5.06°) and chronic 9-16 weeks (172.95° ± 8.88°). Overall, these L5/6 putative inhibitory neurons showed less ability to fire at a consistent phase of intralaminar LFP oscillatory activity during the chronic implantation period. Although the firing rates of detected spikes were stable, the deficits in spike-LFP entrainment and LFP cross-frequency synchronization demonstrate the dysfunctions of the L5/6 intralaminar network near the microelectrode over time.

### 3.3. Imbalance of Cortical Interlaminar Connectivity near the Chronically Implanted Electrode

#### 3.3.1. Direction-dependent impairment in L4 – L2/3 interlaminar connectivity

The chronic implantation injury had been observed to alter the functionality of the intralaminar network in a depth-dependent manner. To further understand the impact of the injury on interlaminar functional connectivity between distinct laminar structures, we first focused on L4, the input cortical layer for visual information from the thalamus, and L2/3, the subsequent processing layer [77, 81]. While our previous findings showed early impairment in the superficial L2/3 with L4 remaining a stable excitatory network over chronic implantation, we now aimed to investigate the changes in functional connectivity between L4 and L2/3.

We first determined the functional connectivity between L4 and L2/3 by analyzing the phase-amplitude coupling (PAC) of theta phase in L4 and gamma amplitude in L2/3. The modulation index (MI) heatmap revealed a reliable PAC relationship between ∼5 Hz phase in L4 and 60-90 Hz amplitude in L2/3 in response to visual stimulation (Figure 5A). Additionally, we assessed the PAC between L4 theta phase and LFP amplitude across all cortical channels. The slow oscillations in L4 were exclusively phase-coupled to gamma frequency amplitude in the superficial layer corresponding to L2/3 (Figure 5B), confirming the functional connectivity between L4 and L2/3 during visual stimulation. However, visual stimulation did not significantly elevate the MI value of L4 phase and L2/3 amplitude compared to the resting state at week 7-10 post-implantation (Figure 5C, *p* = 2.98 * 10^(-11)). This comparable MI value of L4 phase coupling with L2/3 LFP amplitude between visual stimulation and resting state indicated disrupted interlaminar connectivity near the microelectrode. The subsequent increase in MI values since week 11 post-implantation suggested a chronic remodeling of connectivity that enhanced L4-L2/3 synchronization.

**Figure 5.**
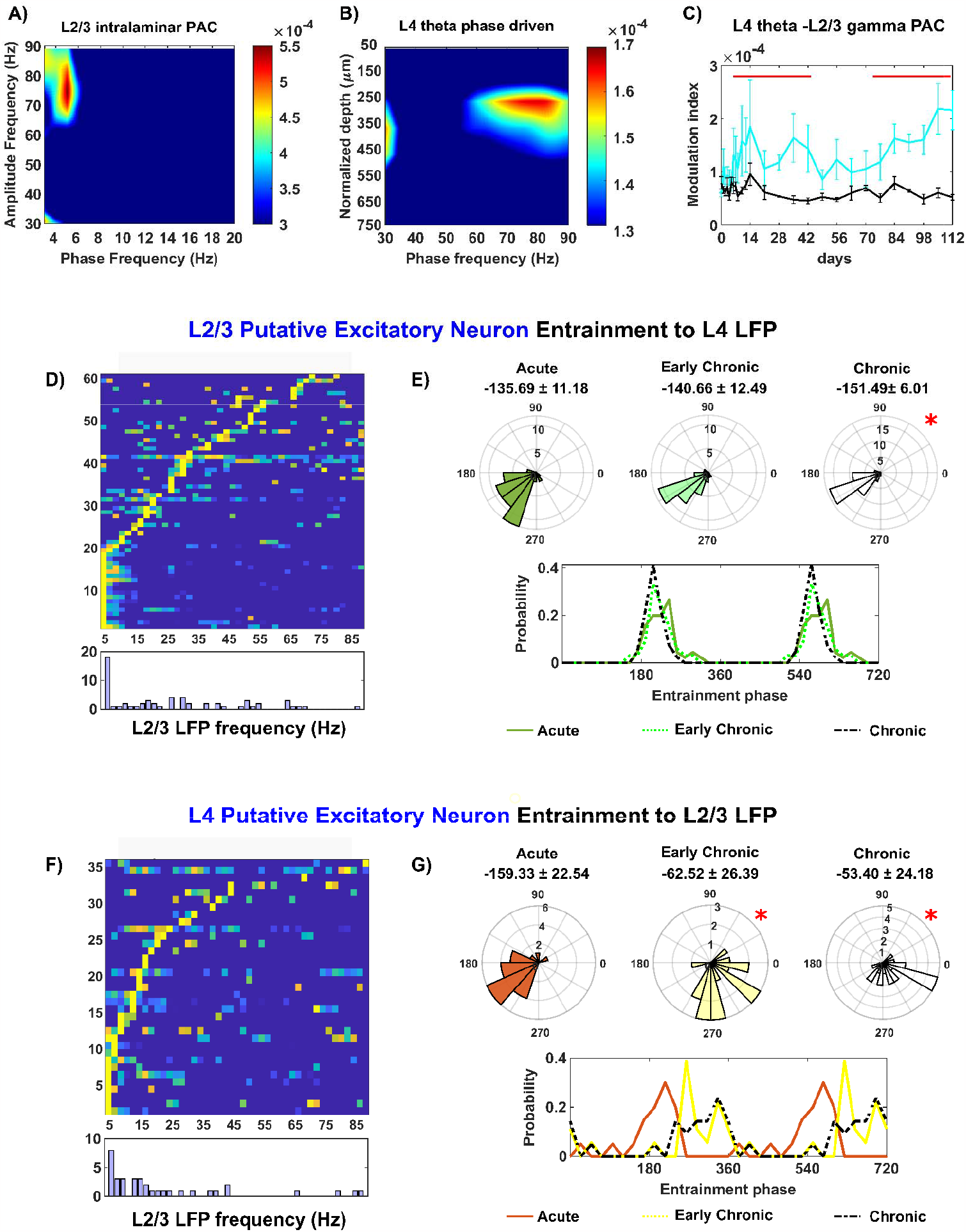
Impaired interlaminar connectivity between L4 and L2/3 during chronic microelectrode implantation. A) Comodulogram displaying the phase-amplitude coupling (PAC) between slow oscillation frequency (2-20 Hz) in L4 and LFP amplitude (30-90 Hz) in L2/3 during visual stimulation. The modulation index (MI) peaked at 5.32 ± 0.15 × 10^-4, indicating strong coupling between 5 Hz theta phase in L4 and 60-90 Hz gamma amplitude in L2/3. B) PAC comodulogram between slow oscillation (4-7.5 Hz theta frequencies) in L4 and LFP amplitude across all cortical channels, highlighting the interlaminar coupling of L4 phase to high gamma amplitude in the superficial layer corresponding to L2/3. C) Plot showing the MI of L4 theta phase-L2/3 gamma amplitude during visual evoked responses (cyan) and resting state (black) over time. D) Interlaminar entrainment of L2/3 putative excitatory neurons to L4 LFP oscillations. The top panel displays the normalized entrainment MI of each L2/3 putative excitatory neuron to L4 oscillations across frequencies, while the bottom panel presents a summary histogram showing that most neurons were phase entrained to 2-7.5 Hz. E) Preferred phase distribution of L2/3 putative excitatory neurons entraining to L4 slow oscillation over different implantation stages, depicted in a polar histogram (top) and a line plot (bottom). F) Preferential entrainment of L4 putative excitatory neurons to L2/3 slow oscillations over the 2-20 Hz frequency range. G) Preferred phase distribution of L4 putative excitatory neurons entraining to L2/3 2-20 Hz oscillation over different implantation stages, represented in a polar histogram (top) and a line plot (bottom). The red bar indicates non-overlapping 95% confidence intervals between visual evoked responses and resting state at each time point, determined using a linear mixed-effects model with likelihood ratio test. The * symbol indicates significant differences in phase angular distribution, either in the resultant mean angles (Watson–Williams test), distribution variability (Kuiper test), or both.

Next, we examined the spike entrainment to LFP oscillations to investigate the directional changes in interlaminar functional connectivity between L2/3 and L4. Recent evidence suggests that neurons exhibit precise firing patterns when receiving organized oscillatory inputs [59, 82, 83]. While inhibitory neurons typically have shorter axons [84], pyramidal neurons, which are the major type of excitatory neurons in the cortex, possess long axons that can span across different laminar structures [85-87]. Making use of our putative subtype classifications in our study, we focused on putative excitatory neurons to examine interlaminar spike-LFP entrainment. For L2/3 putative excitatory neurons, approximately 33% exhibited preferential entrainment to 2-7.5 Hz oscillations in L4 (Fig. 5D). While the mean angles of preferred entrained phase were similar (Watson–Williams test, *p >* 0.05), the variance of the preferred phase distribution was significantly narrowed at the chronic 9-16 weeks (Fig. 5E, Kuiper test < 0.05). This subtle yet significant difference in phase entrainment of L2/3 spikes to L4 oscillatory input indicated changes in interlaminar functional connectivity from L4 to L2/3 near the microelectrode. In reverse direction, 57% putative excitatory neurons in L4 exhibited preferential entrainment to slow 2-20 Hz oscillations in L2/3 (Fig. 3F). However, the phase locking of L4 putative excitatory neurons to L2/3 oscillation underwent substantial changes over chronic implantation, with a significant counterclockwise shift in the mean phase by 100 degrees (Figure 5G, Watson–Williams test, *p <* 0.05). These significant changes in L4 spike entrainment to L2/3 LFP indicated impaired feedback connectivity as L4 excitatory neurons lost their ability to synchronize with L2/3 oscillations. Overall, the chronic implantation injury disrupts the direction-specific functional connectivity between L4 and L2/3, which aligns with the early damage observed in the L2/3 intralaminar network.

#### 3.3.2. Imbalanced L2/3 - L5 mutual connectivity

Although sensory information in L2/3 is typically relayed to the subsequent L5/6 output layer, there is also reciprocal projections from L5 to L2/3 [88-92]. To understand how this mutual connectivity is affected by chronic microelectrode implantation, we examined the interlaminar phase-amplitude coupling (PAC) of LFP oscillations between L2/3 and L5. The comodulogram of L2/3 theta phase revealed strong couplings with LFP amplitude at different cortical depths, including L2/3 itself, L4, and shallow L5 (Fig. 6A). The coupling between L2/3 theta phase and L5 gamma amplitude (∼55-65 Hz) during visual evoked responses was significantly higher than during resting state until week 12 post-implantation (Fig. 6B, *p* = 8.44 * 10^(-9)). However, this coupling progressively declined over time, indicating a loss of L2/3’s ability to synchronize L5 LFP oscillatory activity at the chronic implantation period. In the opposite direction, the L5 theta oscillation was exclusively coupled to ∼60-80 Hz oscillation amplitude in L2/3 (Fig. 6C), indicating the reciprocal functional projection from L5 to L2/3. Interestingly, the coupling between L5 theta and L2/3 gamma during visual evoked responses was consistently higher than during resting state throughout the 16-week implantation period (Fig. 6D, *p* = 1.37 *10(-12)), suggesting an enhanced interlaminar connectivity from L5 to L2/3 near the microelectrode.

**Figure 6.**
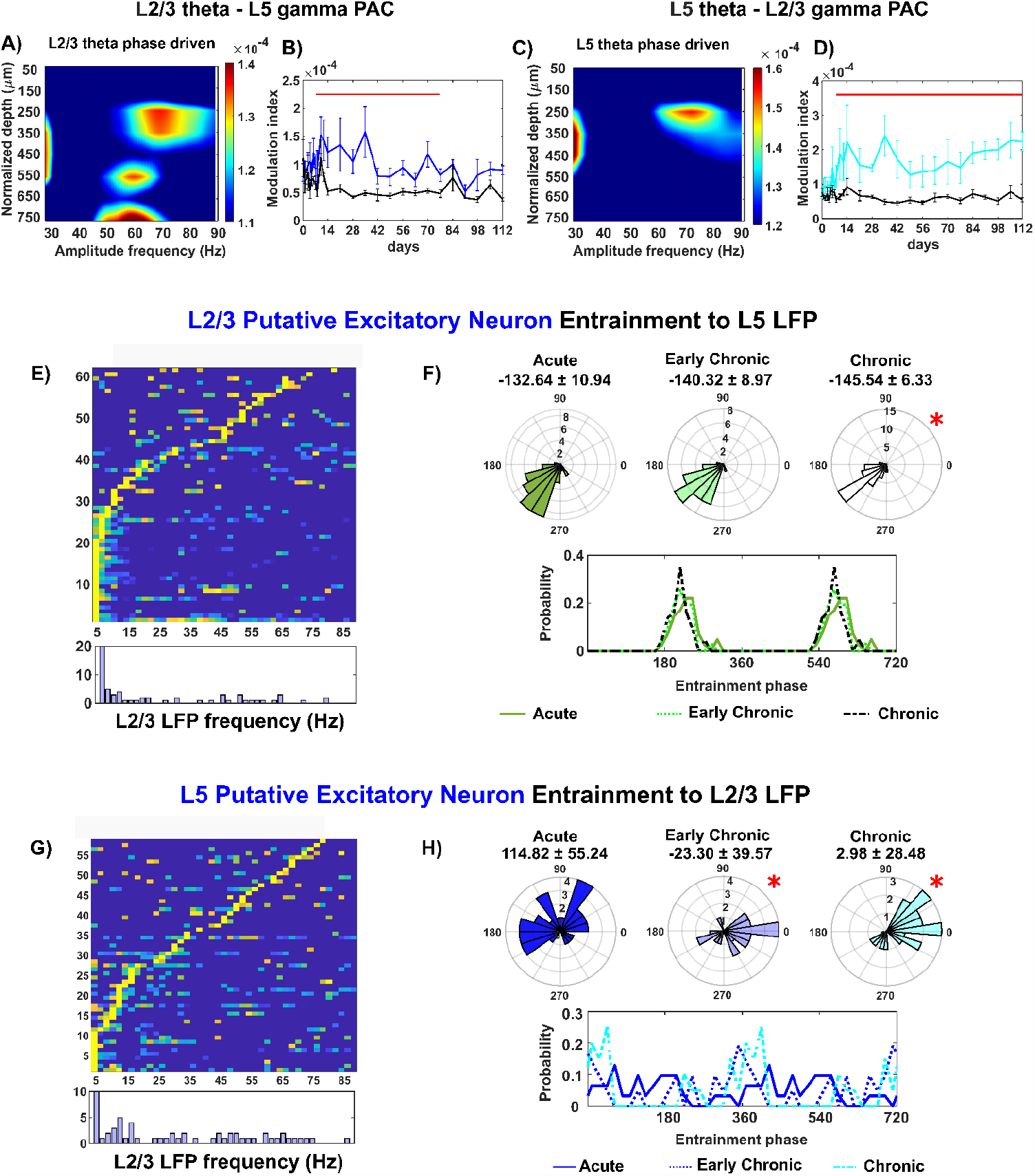
The impaired balance of mutual connectivity between L2/3 and L5 over chronic microelectrode implantation. A). PAC comodulogram between L2/3 theta phases and LFP amplitude across all cortical channels. The robust MI cluster over 55-65 Hz amplitude frequency at 550 μm below surface demonstrates the L2/3-L5 connectivity. B) The MI of L2/3 theta phase coupling to L5 gamma amplitude during visual evoke (blue) gradually decreased to the level of resting state (black) over time. C). The L5 reciprocal projection to L2/3 was represented as the robust L5 theta phase coupling to ∼60-80 Hz amplitude at 250 μm below the brain surface. D). The MI value of this PAC during visual evoke was significantly elevated compared to resting state over the entire 16-week implantation. E). L2/3 putative excitatory neurons were preferentially entrained to L5 2-7.5 Hz oscillation. Top: normalized entrainment MI of each L2/3 putative excitatory neuron to L5 oscillations across frequencies. Bottom: summary histogram. F). Preferred phase distribution of L2/3 putative excitatory neurons entraining to L5 slow oscillation over different implantation stages (Top: polar histogram; bottom: line plot). G). L5 putative excitatory neurons with a Rayleigh test p value < 0.05 were preferentially entrained to 2-20 Hz oscillations in L2/3. H). Preferred phase distribution of L5 putative excitatory neurons entraining to L2/3 2-20 Hz oscillation over different implantation stages (Top: polar histogram; bottom: line plot). Red bar indicates non-overlapping 95% confidence intervals between visual evoke and resting state at each time point using a linear mixed effects model with likelihood ratio test. The * indicates the significant differences between phase angular distribution in either the resultant mean angles (Watson–Williams test) or distribution variability (Kuiper test), or both.

Spike entrainment analysis further revealed changes in the synchronization between L2/3 and L5 neurons to LFP oscillations. For L2/3 putative excitatory neurons, 38% exhibited preferential entrainment to L5 slow 2-7.5 Hz oscillation (Fig. 6E). Additionally, there was a shift in phase locking to L5 slow oscillation in a clockwise direction at the chronic implantation stage (Fig. 6F, Watson–Williams test, *p <* 0.05). Visual observation demonstrated a reduced probability of ∼240° entrained phases at chronic implantation stage. Conversely, nearly 48% of putative excitatory neurons in L5 were entraining to 4-20 Hz oscillation in L2/3 (Fig. 6G). L5 putative excitatory neurons showed a substantial degradation in their synchronization to L2/3 oscillatory input, with significant changes in both mean phases and variance of preferred phase distribution over time (Fig. 6H, Watson–Williams test, *p <* 0.05; Kuiper test, *p <* 0.05). These results indicate a direction-specific impairment in the interlaminar connectivity between L2/3 and L5, leading to imbalanced mutual communication during information processing in the cortex. The dysfunctions observed in L2/3 intralaminar networks likely contribute to the impaired functional projection from L2/3 to L5, while the enhanced functionality of the L5 projection back to L2/3 suggests compensatory mechanisms during chronic implantation.

### 3.4. Disruption of Hippocampal Network Activity during Chronic Microelectrode Implantation

As sensory information is relayed from cortex to hippocampus, we investigated the laminar network near the microelectrode tip in the hippocampal CA1 region. While the cortical network near the implanted microelectrode showed significant activation in response to visual stimuli, the firing rate profile in the hippocampus CA1 region exhibited limited network responses (Fig. 1). To assess the activation of the CA1 network near the microelectrode, we compared the phase-amplitude coupling (PAC) synchronization within CA1 LFP oscillations between visual evoke and resting state conditions. Although the CA1 theta oscillation was coupled to intralaminar gamma amplitude (Fig. 7A, 7B), there was no significant difference in PAC modulation index (MI) between these two conditions over time (Fig. 7C). Furthermore, the firing rates of putative excitatory neurons in CA1 did not show a statistically significant difference between visual evoke and resting state throughout the 16-week implantation (Fig. 7D). These findings suggest that the CA1 circuit near the implanted microelectrodes in this study was not strongly activated during drifting bar visual stimulation. Hippocampus is highly involved in processing visual scenes for discrimination and memory recall [47, 93, 94]. It is possible that a drifting-bar gradient visual stimulus may not provide a strong input to the hippocampal network, or the microelectrode tip may be positioned in a region of CA1 that is less responsive to our visual stimulus paradigm.

**Figure 7.**
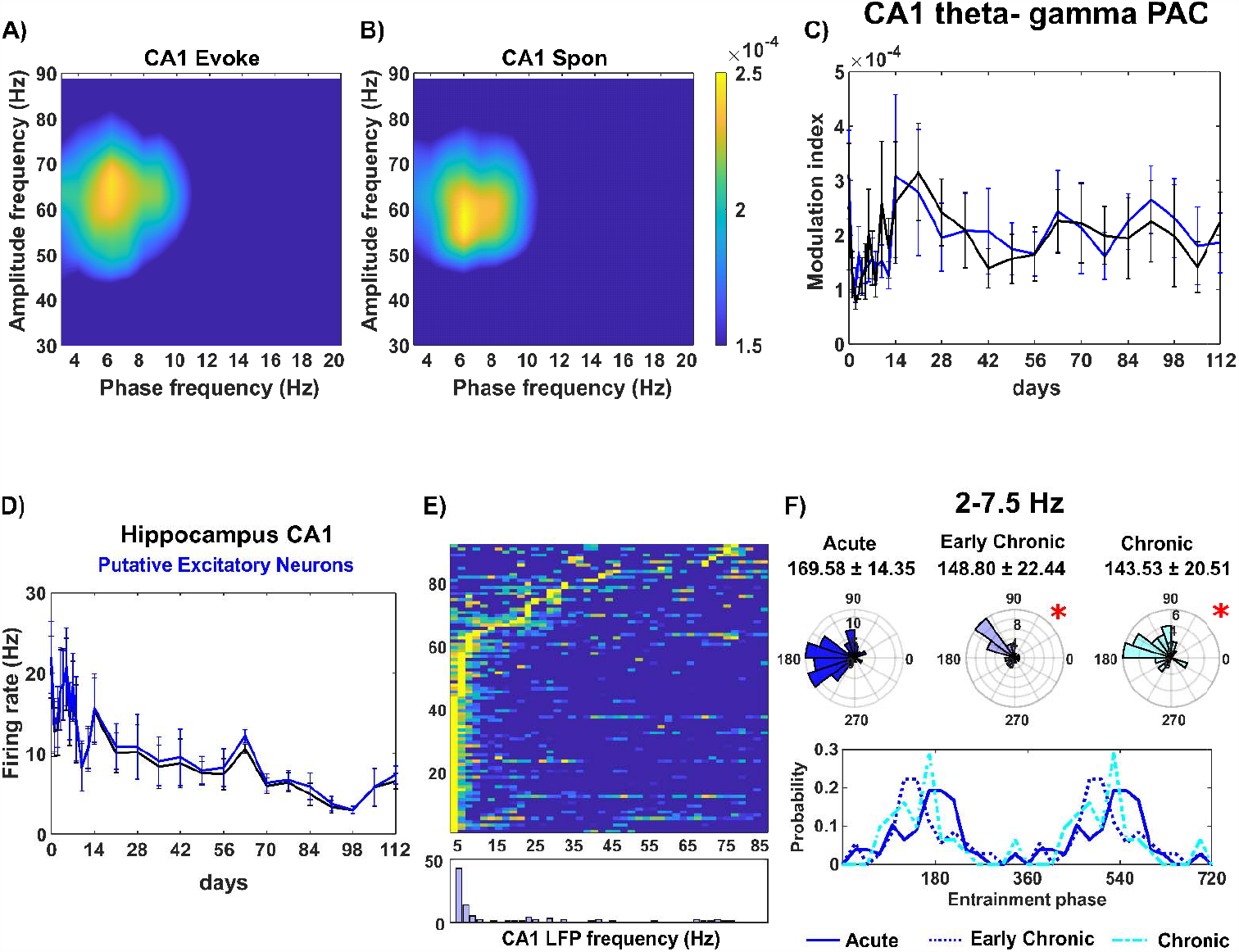
Disruption of hippocampus CA1 communication over chronic implantation. PAC comodulogram of CA1 LFP oscillation between slow frequency (2-20 Hz) phases and high-frequency (30-90 Hz) amplitudes during visual evoke (A) and resting state (B). Robust MI coupling was observed between ∼4-8 Hz phases and 50-70 Hz amplitude in both conditions. C). The MI of CA1 theta-gamma PAC during visual evoke (cyan) was comparable to that during resting state (black) over time. D). Firing rates of putative excitatory neurons in CA1 during visual evoke (blue) and resting state (black) over time. E). The majority of CA1 putative excitatory neurons exhibited preferential entrainment to CA1 2-7.5 Hz oscillation. Top: Normalized entrainment MI of each CA1 putative excitatory neuron to intralaminar LFP oscillations across frequencies. Bottom: Summary histogram. F). Preferred phase distribution of CA1 putative excitatory neurons entraining to 2-7.5 Hz oscillation across different implantation stages (Top: Polar histogr m; Bottom: Line plot). * Indicates significant differences in phase angular distribution, including resultant mean angles (Watson–Williams test) and distribution variability (Kuiper test), or both.

However, microelectrode implantation did result in changes in the hippocampus CA1 network. The firing rate of CA1 putative excitatory neurons peaked at day 7 and then gradually decreased over time (Fig. 7D). Given that pyramidal neurons in CA1 typically have firing rates below 10 Hz [95], the transient peak in firing rate suggests an acute hyper-activity in the network adjacent to the microelectrode. Additionally, the entrainment analysis of CA1 putative excitatory neurons to intralaminar LFP oscillations revealed impairments in local neuron communication. Around 63% of putative excitatory neurons in CA1 exhibited preferential phase entrainment to 2-7.5 Hz slow oscillation (Fig. 7E). The phase locking behavior in the acute 0-2 weeks stage (169.58° ± 14.35°) was significantly different compared to the early chronic 3-8 weeks (148.79° ± 22.44°) and chronic 9-16 weeks (143.53° ± 20.51°) stages (Fig. 7F). The mean phase showed a significant clockwise shift (Watson–Williams test, *p* < 0.05), and the distribution of preferred phases exhibited increased variability (Kuiper test, *p* < 0.05) over time. Visual observation suggests that both early chronic and chronic implantation stages reduced the probability of preferred phases ∼180°-240° compared to the acute 0-2 weeks stage. These results indicate that the CA1 neurons adjacent to the microelectrode were less able to respond appropriately to synchronized oscillatory inputs. In summary, chronic implantation led to local impairments in the CA1 circuit, including a chronic dysfunction in individual neural response to surrounding LFP oscillations.

### 3.5. Histology of Neuronal Subtypes in Different Brain Laminar Structures Near the Chronic Microelectrode

Given electrophysiological changes in functional network connectivity over time, we assessed the integrity of neuronal populations following 16-week chronic implantation by immunohistology. Immunohistochemical staining markers were used to label different neuronal subtypes (CaMKIIα for excitatory neurons and GAD67 for inhibitory neurons) and dendrites labeled by MAP2. Coronal sections of the tissue were examined to visualize the depth distribution of these markers in the cortex and hippocampus CA1.

In the cortex, a reduction in fluorescence intensity of CaMKIIα (Fig. 8A) and GAD67 (Fig. 8B) was observed near the implant site, particularly in the superficial cortical layers. This loss indicated damage to both excitatory and inhibitory neurons in the superficial cortical depth. Additionally, a significant decrease in MAP2 fluorescence was observed at superficial depths (Fig. 3C), suggesting impaired dendritic structure in the superficial network due to the implantation injury. These histological findings corresponded with the electrophysiological measurements showing impairment in the superficial L2/3 region of the cortex.

**Figure. 8.**
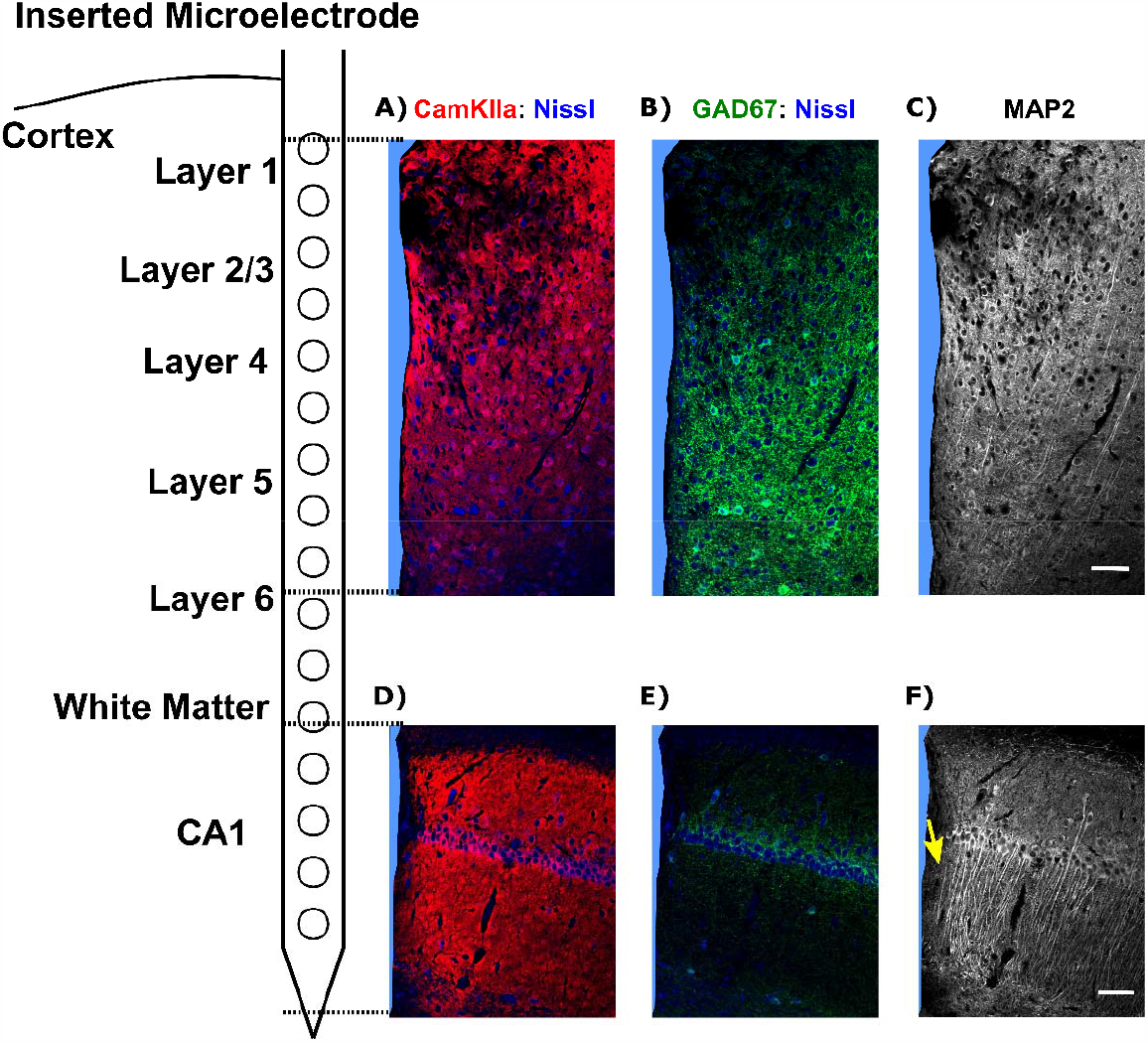
Histological evaluation reveals prominent loss of neural markers at superficial cortical depth relative to deep structures. Representative histological images demonstrated loss of CaMKII α+ Nissl+ excitatory neurons (A), GAD67+ Nissl+ inhibitory neurons (B), and MAP2+ dendrites (C) near the microelectrode at superficial cortical depths. Images of the hippocampus CA1 showed the distribution of CaMKII α+ Nissl+ excitatory neurons (D), GAD67+ Nissl+ inhibitory neurons (E), and MAP2+ dendrites (F) near the chronically implanted microelectrode. Microelectrode shanks are labeled in blue. Scale bar = 50 μm.

In the hippocampal CA1 region, dense CaMKIIα+ Nissl+ soma showed reduced fluorescence intensity adjacent to the implant (Fig. 8D), indicating the impairment of CA1 pyramidal neurons located in the stratum pyramidal (SP). Few GAD67+ Nissl+ cells were observed in the hippocampal CA1 region, making it difficult to assess inhibitory network integrity in this area (Fig. 8E). Interestingly, low levels of MAP2 fluorescence were observed in the region beneath the CA1 soma layer (Fig. 8F, yellow arrow), corresponding to the CA1 stratum radiatum (SR) where dendrites input to pyramidal cells in the SP. This finding suggested that chronic microelectrode implantation impaired the appropriate synaptic inputs to pyramidal neurons. Overall, these histological observations supported the electrophysiological results, demonstrating depth-dependent tissue changes caused by chronic microelectrode implantation.

## 4. Discussion

Intracortical microelectrodes have demonstrated potential as a therapeutic tool for neurological disorders [3, 4, 10, 13, 96]. However, the gradual decline in recording quality during long-term implantation leads to the instability of BCI decoders over time creating a major challenge in their clinical application [14, 24, 57, 97]. BCI decoders are algorithms that translate brain signals into digital commands for computer and robotic outputs [98]. However, the brain signals generated by the neural networks are subject to changes produced by tissue degeneration such as neuronal density loss and glial encapsulation near the implant site [28]. The instability of brain signals can degrade BCI performance, requiring frequent recalibration of the BCI decoders [13, 99, 100], a time- and effort-consuming process that causes substantial inconvenience to the patients during each session.

To facilitate addressing these challenges, this study aims to examine how functional network connectivity near the implanted microelectrode changes over time. Our findings revealed a depth-dependent degradation in intralaminar functional connectivity. The superficial layers (L2/3) showed early impairment with a loss of putative inhibitory neuron detection and a significant decrease in the firing rate of putative excitatory neurons. Interlaminar functional connectivity changes near the implant were found to be directional, specifically between L2/3 and L5. In the hippocampal CA1 region near the implant there were changes in the precise timing of spikes in response to oscillatory local field potential (LFP) activity. These findings provide valuable insights into the changes in functional network connectivity due to tissue reactions near the chronically implanted microelectrode, separate from neuroplasticity changes linked to learning. Elucidating the mechanisms behind the decline in signal quality over time can contribute to the design of therapeutic strategies for improving the tissue health near the chronically implanted electrode. Additionally, this knowledge can inform the development of more stable and reliable BCI decoders.

### 4.1. Changes to excitatory activity may contribute to dysfunctions in laminar network connectivity near the chronically implanted microelectrode

The majority of cortical neurons are excitatory glutamatergic neurons, interconnected within and across layers to facilitate recurrent excitation in brain circuits [101]. This local recurrent excitation could implement amplification and thus elevate network connectivity [102]. However, networks with high amplification are sensitive to disruption, since the loss of even a few neurons involved in recurrent excitation can impair information processing in the L2/3 network [101]. Our findings reveal a layer-dependent profile of putative excitatory neuron firing rates which decreased significantly in L2/3 during the chronic implantation period (Fig. 3D). This observation suggests a disruption in recurrent excitation within the L2/3 local network, which may affect L2/3 network connectivity. Additionally, L2/3 putative excitatory neurons exhibit improper entrainment to intralaminar LFP oscillatory input (Fig. 3E-G) during the chronic implantation period, providing evidence of dysfunction in excitatory activity in the L2/3 intralaminar network. The recurrent network of excitatory neurons in L2/3 plays a vital role in sensory processing including feature extraction, specific input amplification, and information integration across different sensory modalities [86, 101, 103]. Therefore, impairment to L2/3 excitatory activity near the chronically implanted microelectrode could result in network functionality deficits, potentially affecting the quality and reliability of signals received from the implant for further BCI applications.

Moreover, L2/3 serves as a crucial relay station for ascending and descending projections, mediating communication between superficial and deep cortical layers [86]. Recurrent excitation exists between L2/3 and L5 as L2/3 pyramidal neurons play a vital role in amplifying sensory-evoked responses in L5 neurons [81]. Therefore, the impairment of L2/3 excitatory activity may affect its interlaminar excitatory projection to L5, which could explain the substantial degradation of L5 putative excitatory neuron synchronization to L2/3 oscillatory input (Fig. 6G-H). Furthermore, the excitatory activity within the L5/6 intralaminar network is influenced, as suggested by the changes in responsiveness of the L5/6 putative excitatory neurons to intralaminar LFP oscillatory activity over time (Fig.4E-G). Thus, disruption in the excitatory activity of L2/3 may be associated with the disturbances in the deeper layers that serve as cortical outputs near the chronically implanted microelectrode. However, further studies are necessary to ascertain whether the changes in L5/6 putative excitatory neuron activity result from a cascading impact originating from L2/3 disruptions, or whether they are directly influenced by chronic implantation injury. Potential future investigations could focus on unraveling the sequence of these changes to deepen our understanding of chronic alterations in cortical networks near the implanted microelectrode. One possible approach could involve confining the microelectrode implantation to L2/3 and scrutinizing excitatory activity through the assessment of glutamate synaptic density and postsynaptic calcium dynamics in the deep layers L5/6. This evaluation may help examine whether there is a cascade effect from L2/3 to L5/6 when the microelectrode does not cause injury in the deep layers.

In addition to cortical networks, the hippocampus CA1 region also experiences a shift in the entrainment of putative excitatory neurons to LFP oscillatory input (Fig. 7), suggesting a disruption in appropriate excitatory synaptic connectivity in CA1 near the implanted microelectrode. The stiff microelectrode implantation is known to cause more concentrated mechanical strain in the deeper tissue near the tip, as compared to the superficial layers [25]. In this study, microelectrodes were perpendicularly inserted 1600 μm below the brain surface, positioning the tip near the CA1. Thus, brain tissue in CA1 near the tip is likely to experience increased mechanical strain, potentially triggering microglial mechanosensory response and leading more microglial cells to transform to an activated state [104]. Implantation-induced inflammation has been associated with the activation of Toll-like receptor (TLR) 2 and TLR4 in microglia, which can reduce long-term potentiation (LTP) [105]. This could be a potential pathway leading to the observed disruption of putative excitatory neurons’ response to LFP synaptic input in CA1 (Fig. 7F). However, the comprehensive mechanisms through which inflammation impacts synaptic functionality in the neural circuits near the implanted microelectrode remain to be fully elucidated and warrant further research.

### 4.2. Changes to inhibitory activity may contribute to dysfunctions in laminar network connectivity near the chronically implanted microelectrode

Although representing only a small fraction of the total neuronal population, inhibitory neurons play a critical role within neural networks [69, 106]. Inhibitory neuron activity helps to balance excitation in the network, and this balance is essential for proper information processing in the brain [78, 107]. Feedback inhibition is a common activity motif at the network level, involving a set of excitatory neurons stimulating inhibitory neurons, which in turn suppress the activity of the same population of excitatory cells [108, 109]. This feedback inhibition is vital for synchronization of populational firing of excitatory neurons [108]. In this study, we noted a decline in the involvement of L2/3 putative inhibitory neurons in visually evoked activation near the implanted microelectrode (Fig. 3C), suggesting an early impairment of L2/3 intralaminar feedback inhibition. Additionally, alterations in L2/3 intralaminar spike entrainment to LFP (Fig. 3E-J), along with decreased firing rates of putative excitatory neurons (Fig. 3D), indicate a loss of synchronized excitatory activity. These observations collectively suggest a disruption in the excitation-inhibition balance in the L2/3 near the microelectrode, which may influence accurate information processing. However, additional research is needed to investigate whether these modifications in L2/3 inhibitory neuronal activity are the driving force behind changes in excitatory responses to chronic implantation injury.

Moreover, impairment in inhibitory activity could potentially disrupt interlaminar communication with deeper layers. Feed-forward inhibition typically occurs between different cortical layers, where excitatory neurons stimulate intralaminar inhibitory cells that subsequently inhibit a group of postsynaptic excitatory neurons in another layer [108, 110]. Thus, feed-forward inhibition is critical for maintaining interlaminar network connectivity [108, 110]. Our observations indicate a loss of functional network connectivity from L2/3 to L5 (Fig. 6A-B) and from L2/3 to L4 (Fig. 5F-G), along with the loss detection of L2/3 putative inhibitory neurons (Fig. 3). These findings suggest that the impairment of inhibitory activity in L2/3 may lead to deficits in feed-forward inhibition to the deeper layers, explaining the decreased interlaminar network connectivity with L2/3 in the descending direction. Furthermore, it is likely that diminished inhibitory input from L2/3 affects the balance of excitation and inhibition within the L5 intralaminar network, subsequently contributing to alterations in synchronization from L5 to L2/3. Interestingly, we noted that the connectivity in the reverse ascending direction, from L4 to L2/3 (Fig. 5C) and L5 to L2/3 (Fig. 6C-D), is heightened. This heightened interlaminar connectivity from L5 to L2/3 could potentially indicate abnormal amplification from L5 due to a loss of L2/3 feed-forward inhibition. An imbalance in the reciprocal connectivity between L2/3 and L5 could thus alter network dynamics and result in modifications to neuronal entrainments to LFP oscillatory input. Future studies are required to determine if, and how, the impairments in inhibitory activity in L2/3 directly contribute to the observed changes in the interlaminar connectivity with deeper layers. Additionally, the specific impact of these changes on the excitation-inhibition balance within L5 intralaminar network and on network dynamics should be explored.

### 4.3. Changes in metabolic activity may contribute to alterations in laminar network connectivity near the microelectrode

Deficits in metabolic supply to neurons may contribute to dysfunctions of neural networks. The observed impairments in the network functionality of superficial layers 2/3 (Fig. 3, Fig 8) could be linked to vascular injuries induced by electrode implantation, resulting in more severe tissue damage in superficial depths with large-diameter vessels than in deeper tissues with smaller capillaries [111-113]. Additionally, astrocytes [35, 114, 115] and oligodendrocytes [41, 43] that mediate local delivery of metabolites to neurons undergo more structural changes in response to implantation injury at superficial layers [116, 117], which may further compromise the metabolic supply for neural network functionality near the microelectrode. Therefore, this disruption to vascular integrity and glial reactivity potentially reduces the availability of glucose and oxygen for neuronal activity [28, 31, 40], which may explain the early detection loss of metabolically intensive inhibitory neurons, significant reduction in putative excitatory firing rate, and decreased network activation in L2/3 (Fig. 3). Further research is needed to provide more direct evidence elucidating the relationship between glial activity and vasculature dynamics in the inflammatory environment and deficits in neural network functionality from a metabolic perspective.

Inefficient waste removal may contribute to changes in laminar network connectivity near the chronically implanted microelectrode. A buildup of toxic substances to neurons resulted from neuroinflammation following electrode implantation may be associated with the progressive degeneration of L2/3 network functionality (Fig. 3). While microglia with phagocytic capacity quickly become activated in response to microelectrode implantation, there is a mismatch between this immediate microglial activity and the subsequent neuron loss near the microelectrode [118-122]. Microglial phagocytic repair might not effectively remove the cell debris, leading to sustained neuroinflammation, particularly in the superficial layers. Future studies could explore the impacts of implantation on microglial phagocytosis and how these might influence the functionality of the implanted microelectrode.

### 4.4. Network instability and decoder performance: decoupling plastic learning and network degeneration in disease and injury

The stability of BCI decoders, which convert brain signals into digital information, is crucial for their effective performance, particularly in long-term applications like neuroprosthetics and clinical treatments. However, the long-term decoder performance is hindered by the neural reorganization produced by implantation-induced injury and/or “plasticity” [13, 99, 100]. Therefore, gaining a comprehensive understanding of changes in neural network activity can assist in designing decoders with machine learning algorithms that can adapt to shifting neural data and maintain their performance over time. In this study, we provide a unique perspective on local disturbances in both intra- and inter-laminar network connectivity in response to long-term implantation injury. Our data reveals a gradual decline in single-unit firing rates and alterations in spike entrainment to local field potentials (LFP), indicating dysfunctions in the local neural network near the implanted microelectrode. Previous evidence has shown that these local changes, affecting specific subsets of neurons, directly contribute to persistent deficits in the efficient control of movement over time [18]. Taken together, these findings suggest a potential association between implantation-induced network changes and instability in BCI performance.

In BCI field, the specific mechanisms underlying changes in neural networks during long-term implantation injury compared to neuroplasticity associated learning remain an open question. Previous studies have indicated that learning can occur at multiple stages within the feedback control loop [18]. In the context of a brain-computer interface (BCI) learning paradigm, changes in neural activity patterns are necessary for neurons recorded from implanted microelectrodes to facilitate learning and improve output behavior. In our study, we observed longitudinal changes in the local network surrounding the implanted microelectrode over a period of 16 weeks. If implantation injury causes neuronal damage within the learning network, it may diminish the efficiency of learning and contribute to BCI decoder instability over time. However, it is unclear whether the reorganization of neural activity recorded by the BCI device primarily results from neuronal impairment due to implantation injury or alterations induced by learning. Further research is needed to investigate the extent to which neural reorganization near the implanted microelectrode is driven by injury or learning, providing valuable insights into the mechanisms underlying decoder instability.

### 4.5. Limitations

There are several limitations in this study. First, the classification of putative neuronal subtypes based on spike width has some constraints. While this approach is commonly used in many studies to separate sorted waveforms into different functional groups [58, 126], it is important to note that further validation is required to confirm the physiological identification of these neuronal subtypes [127]. Optogenetic strategies, combined with electrophysiological measurements, could be applied to validate the identification of neurons classified based on spike width. Alternatively, other parameters such as firing rates, depth location, and monosynaptic connections through short-latency spike cross-correlograms could be used to further cluster functional subtypes of sorted single-unit waveforms. Moreover, additional immunohistological quantifications would be helpful to understand the longitudinal impact of implantation injury on different brain laminar structures. While CaMKIIa is widely used for labeling excitatory neurons, multiple types of GABAergic interneruons are also labeled by the viral expression driven by CaMKIIa promotor [128]. Therefore, markers such as VGLUT could be included to determine the population of excitatory neurons near the implanted microelectrode. However, it should be acknowledged that despite the limitations, our results still provide insights into different neuronal subtypes with distinct electrophysiological features in response to chronic implantation injury.

Secondly, we utilized cross-frequency coupling (PAC) and spike-LFP coupling to investigate functional connectivity within and across different laminar structures. However, whether the changes in functional connectivity involve changes in anatomical network connections in response to microelectrode implantation over chronic time scales remain unknown. While electrophysiological measures provide insights into synaptic and firing activity near the implant, further investigation through anatomical evaluation, such as immunohistochemical staining of presynaptic markers (e.g., Neurexin-3α, SNAP-25, Synapsin) and postsynaptic markers (e.g., Gephyrin, Homer1, Synapse-Associated Protein-102), can provide direct evidence on how synaptic connectivity plasticity is affected near the implant in various laminar structures. These experiments will help bridge the gap between circuit functionality and tissue reactions, providing a deeper understanding of the underlying mechanisms.

A third limitation inherently arises from the sampling of the electrophysiological activity from a random set of neurons surrounding the microelectrode. The limits of a recording site allow for the detection of only about 5 single units [129], yet the neuronal density near the implant is about to 1500 cells/mm^2^ at 50-100 μm [40]. Thus, it becomes challenging to track the same population of neurons over time. Therefore, future studies should consider incorporating in vivo imaging techniques that can reveal the real-time dynamics of neuronal activity. These imaging methods can be performed in parallel with electrophysiological recording during visual stimulation, allowing for a comprehensive assessment of network functionality over chronic time periods. This integrated approach would provide valuable insights into the spatial and temporal characteristics of neuronal activity in response to implantation injury, further enhancing our understanding of the underlying mechanisms and dynamics within the neural circuitry.

## 5. Conclusion

Clinical applications of intracortical microelectrodes face limitations due to decoder instability caused by plastic learning and network degradation resulting from implantation injury. The chronic instability of the recorded signal’s functional network connectivity poses a significant challenge, including the need for regular retraining of the decoder, which consumes substantial time and can be burdensome for patients. To comprehensively understand the impact of implantation injury-related functional degradation, it is crucial to investigate the biological reactions near the chronically implanted microelectrode in conjunction with the degeneration of the functional recording network. In our study, we provide a detailed characterization of changes in functional network activity and synchronization within and across different laminar structures in the cortex and hippocampus. We observed depth-dependent alterations in intralaminar network connectivity with impaired network functionality observed at superficial depths. Furthermore, interlaminar connectivity exhibited direction-dependent changes with reduced connectivity from L2/3 to deeper layers and increased connectivity from L5/6 to superficial depths. These findings contribute to our understanding of the functional responses of the neural network near the microelectrode and shed light on the underlying mechanisms responsible for chronic signal instability and the decline of implanted microelectrodes. By illuminating the intricate dynamics of the functional network and its changes over time, our study paves the way for the design and implementation of more effective interventions to improve the long-term stability and performance of intracortical microelectrodes.

## 6. Acknowledgement

This work was supported by: NIH R01NS094396, NIH R01NS105691, NIH R01NS115707, NIH R03AG072218, NIH R01NS129632 and NSF CBET CAREER 1943906.

---

**Supplementary Table 1.**
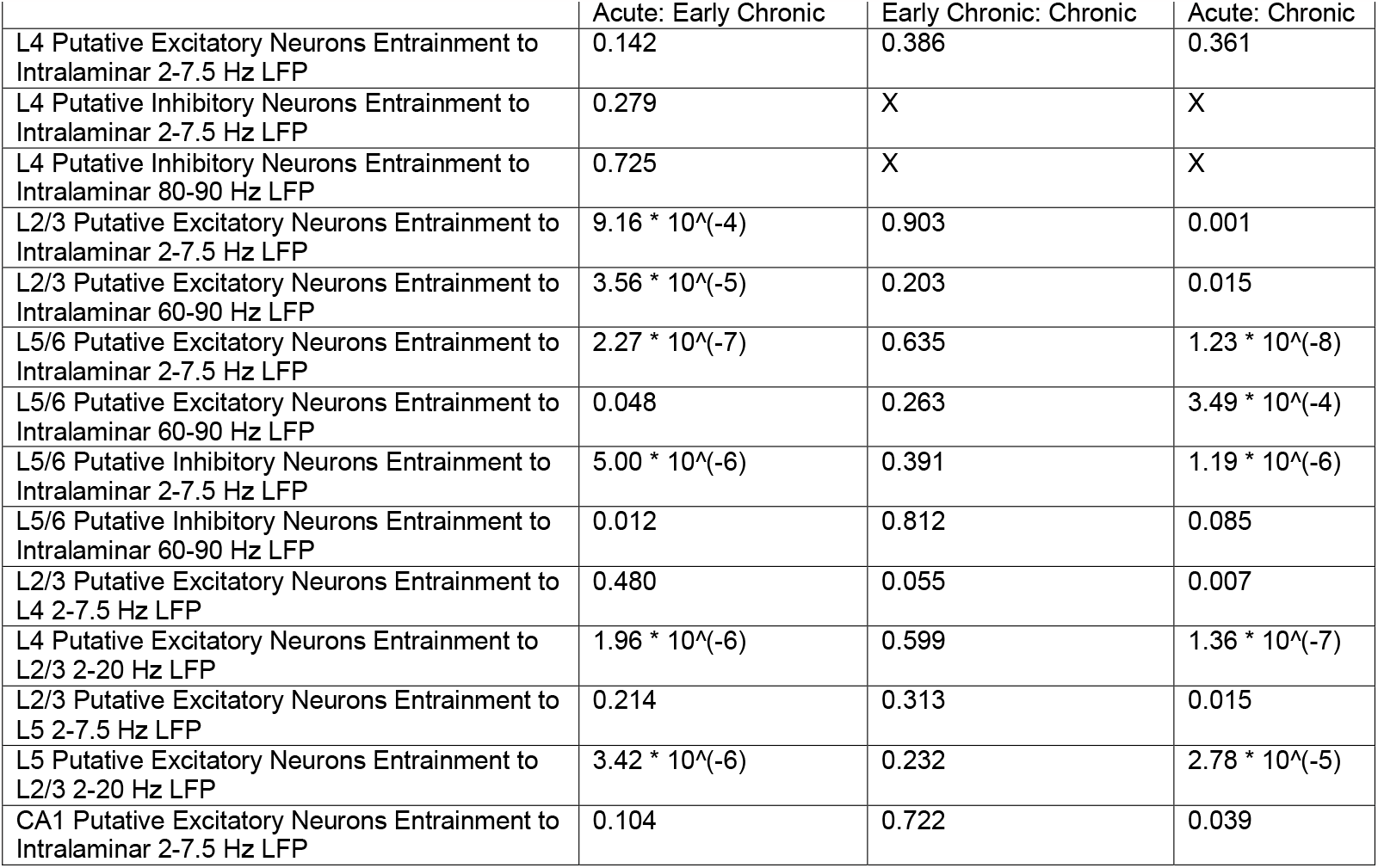
Statistical analysis of Watson-Williams test for the resultant mean entrainment angles; significance *p*<0.05.

**Supplementary Table 2.**
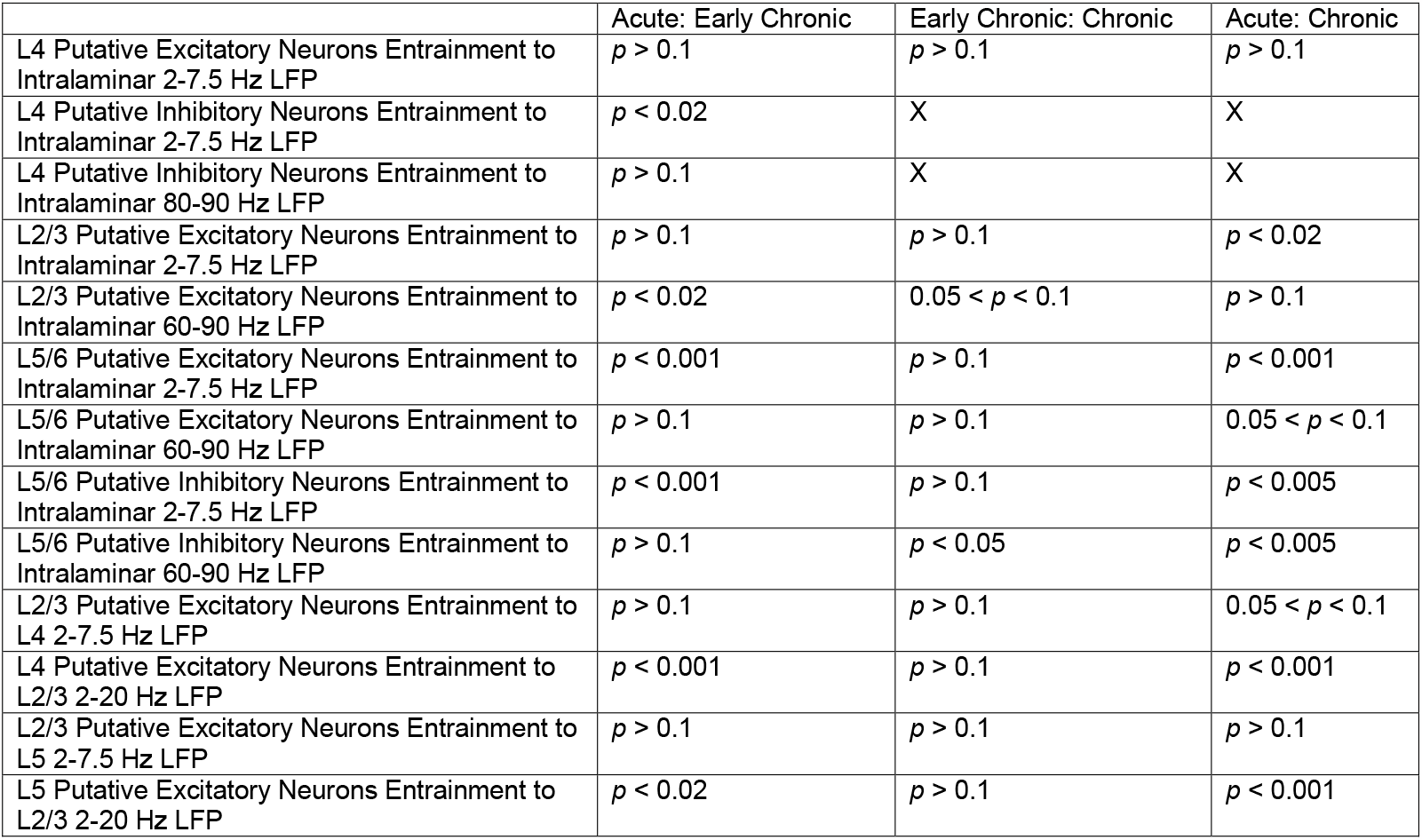
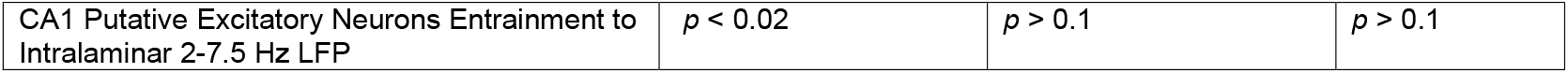
Statistical analysis of Kuiper test distribution of entrainment angles; significance *p*<0.05.

