## Supplemental Tables 1-2 for "Neuronal functional connectivity is impaired in a layer dependent manner near the chronically implanted microelectrodes"

**Keywords:** brain-computer interface, brain circuit, LFP synchronization, visual cortex, neuroinflammation.

**Supplemental Materials**

|  | Acute: Early Chronic | Early Chronic: Chronic | Acute: Chronic |
| --- | --- | --- | --- |
| L4 Putative Excitatory Neurons Entrainment to Intralaminar 2-7.5 Hz LFP | 0.142 | 0.386 | 0.361 |
| L4 Putative Inhibitory Neurons Entrainment to Intralaminar 2-7.5 Hz LFP | 0.279 | X | X |
| L4 Putative Inhibitory Neurons Entrainment to Intralaminar 80-90 Hz LFP | 0.725 | X | X |
| L2/3 Putative Excitatory Neurons Entrainment to Intralaminar 2-7.5 Hz LFP | 9.16 * 10^(-4) | 0.903 | 0.001 |
| L2/3 Putative Excitatory Neurons Entrainment to Intralaminar 60-90 Hz LFP | 3.56 * 10^(-5) | 0.203 | 0.015 |
| L5/6 Putative Excitatory Neurons Entrainment to Intralaminar 2-7.5 Hz LFP | 2.27 * 10^(-7) | 0.635 | 1.23 * 10^(-8) |
| L5/6 Putative Excitatory Neurons Entrainment to Intralaminar 60-90 Hz LFP | 0.048 | 0.263 | 3.49 * 10^(-4) |
| L5/6 Putative Inhibitory Neurons Entrainment to Intralaminar 2-7.5 Hz LFP | 5.00 * 10^(-6) | 0.391 | 1.19 * 10^(-6) |
| L5/6 Putative Inhibitory Neurons Entrainment to Intralaminar 60-90 Hz LFP | 0.012 | 0.812 | 0.085 |
| L2/3 Putative Excitatory Neurons Entrainment to L4 2-7.5 Hz LFP | 0.480 | 0.055 | 0.007 |
| L4 Putative Excitatory Neurons Entrainment to L2/3 2-20 Hz LFP | 1.96 * 10^(-6) | 0.599 | 1.36 * 10^(-7) |
| L2/3 Putative Excitatory Neurons Entrainment to L5 2-7.5 Hz LFP | 0.214 | 0.313 | 0.015 |
| L5 Putative Excitatory Neurons Entrainment to L2/3 2-20 Hz LFP | 3.42 * 10^(-6) | 0.232 | 2.78 * 10^(-5) |
| CA1 Putative Excitatory Neurons Entrainment to Intralaminar 2-7.5 Hz LFP | 0.104 | 0.722 | 0.039 |

**Supplementary Table 1**. Statistical analysis of Watson-Williams test for the resultant mean entrainment angles; significance *p*<0.05.

|  | Acute: Early Chronic | Early Chronic: Chronic | Acute: Chronic |
| --- | --- | --- | --- |
| L4 Putative Excitatory Neurons Entrainment to Intralaminar 2-7.5 Hz LFP | *p* > 0.1 | *p* > 0.1 | *p* > 0.1 |
| L4 Putative Inhibitory Neurons Entrainment to Intralaminar 2-7.5 Hz LFP | *p* < 0.02 | X | X |
| L4 Putative Inhibitory Neurons Entrainment to Intralaminar 80-90 Hz LFP | *p* > 0.1 | X | X |
| L2/3 Putative Excitatory Neurons Entrainment to Intralaminar 2-7.5 Hz LFP | *p* > 0.1 | *p* > 0.1 | *p* < 0.02 |
| L2/3 Putative Excitatory Neurons Entrainment to Intralaminar 60-90 Hz LFP | *p* < 0.02 | 0.05 < *p* < 0.1 | *p* > 0.1 |
| L5/6 Putative Excitatory Neurons Entrainment to Intralaminar 2-7.5 Hz LFP | *p* < 0.001 | *p* > 0.1 | *p* < 0.001 |
| L5/6 Putative Excitatory Neurons Entrainment to Intralaminar 60-90 Hz LFP | *p* > 0.1 | *p* > 0.1 | 0.05 < *p* < 0.1 |
| L5/6 Putative Inhibitory Neurons Entrainment to Intralaminar 2-7.5 Hz LFP | *p* < 0.001 | *p* > 0.1 | *p* < 0.005 |
| L5/6 Putative Inhibitory Neurons Entrainment to Intralaminar 60-90 Hz LFP | *p* > 0.1 | *p* < 0.05 | *p* < 0.005 |
| L2/3 Putative Excitatory Neurons Entrainment to L4 2-7.5 Hz LFP | *p* > 0.1 | *p* > 0.1 | 0.05 < *p* < 0.1 |
| L4 Putative Excitatory Neurons Entrainment to L2/3 2-20 Hz LFP | *p* < 0.001 | *p* > 0.1 | *p* < 0.001 |
| L2/3 Putative Excitatory Neurons Entrainment to L5 2-7.5 Hz LFP | *p* > 0.1 | *p* > 0.1 | *p* > 0.1 |
| L5 Putative Excitatory Neurons Entrainment to L2/3 2-20 Hz LFP | *p* < 0.02 | *p* > 0.1 | *p* < 0.001 |
| CA1 Putative Excitatory Neurons Entrainment to Intralaminar 2-7.5 Hz LFP | *p* < 0.02 | *p* > 0.1 | *p* > 0.1 |

**Supplementary Table** **2**. Statistical analysis of Kuiper test distribution of entrainment angles; significance *p*<0.05.
